# Genome-wide structural variant landscape following HDR-enhanced CRISPR editing in human hematopoietic stem and progenitor cells

**DOI:** 10.64898/2026.06.13.732078

**Authors:** Fanny-Meï Cloarec-Ung, Guy Sauvageau, Hilary M Sheppard, Peter M. Lansdorp, David JHF Knapp

## Abstract

DNA-PK inhibitors (DNA-PKi) have emerged as potent enhancers of homology-directed repair (HDR) in genome editing, however, recent studies have reported that such inhibitors can induce large-scale structural variations (SVs) raising safety concerns. To investigate factors influencing SV induction in human hematopoietic stem and progenitor cells (HSPCs), we employed a combination of on-target long-read nanopore sequencing and unbiased genome wide detection of SVs using Strand-seq. Our results show that editing alone results in a high frequency of large on-target deletions which were modestly increased by DNA-PK inhibition and could be mitigated using DNA polQ inhibitors. Genome-wide SVs were also induced at low frequencies by editing but were not substantially increased with DNA-PKi. SVs including translocations were, however, increased with multiple simultaneous editing. Our results highlight that edit-induced SVs are not limited to DNA-PKi enhanced protocols, thus careful assessment will be critical to ensure the safety of any edited therapeutic cell product.

## Introduction

Genome editing in human hematopoietic stem and progenitor cells (HSPCs) is challenging due to their quiescence and limited maintenance under ex vivo culture conditions(1). In recent years, several studies have demonstrated that the addition of DNA-PK inhibitors, including the molecule AZD7648, can enhance homology-directed repair (HDR)-mediated CRISPR-Cas9 genome editing in both cell lines and primary cells(2–7). Using the inhibitor AZD7648, we and others have achieved high HDR efficiencies across different loci and protocol designs, confirming AZD7648 as a strong strategy for precise genome editing(3–5,8,9). Mechanistically, HDR-mediated CRISPR editing relies on homology-directed repair pathways. The predominant path for repairing double stranded breaks, however, is non-homologous end joining (NHEJ). NHEJ depends on the recruitment of the Ku70/Ku80 complex, which in turn activates DNA-PKcs as a catalytic subunit(10). DNA-PK inhibitors, such as AZD7648, act at this stage to suppress NHEJ, thus requiring a cell to repair using either microhomology-mediated end joining (MMEJ), theta-mediated end joining (TMEJ), a distinct PolQ-dependent pathway, or HDR(11–13). While MMEJ and TMEJ are often considered the same, they rely on different molecular machinery and can produce distinct repair outcomes, a distinction with implications for the large deletions observed following CRISPR editing(14).

While surviving cells following HDR-mediated editing in the presence of AZD7648 show high efficiencies of correct edits at the targeted site(4–6), this raises the possibility of mis-repair at other sites. We previously demonstrated that AZD7648 had negligible effects on the function of the high self-renewing fractions, however, committed progenitors showed some functional deficit(5). Although our analysis found no increase of small indels at predicted off-target sites, we did not assess large-scale structural variations (SVs) at any other sites(5). Recently, Boutin et al showed that loss-of-heterozygosity (LoH) events occurred at the telomeric region next to the targeted sites which were increased slightly in the presence of AZD7648 (3.9 vs 1.8 % at 7 days), though these levels decreased over time(15), suggesting that clones bearing them were competitively disadvantaged. Similarly, Cullot et al., demonstrated that while AZD7648 significantly enhanced HDR efficiency, its use was associated with the induction of a substantial fraction of large on-target deletions(16). These effects were observed across multiple loci and cell types, including p53-proficient and p53-null cells, primary HSPCs and organoid models. The effect was particularly pronounced in p53-null cells, where up to 40% of large deletions were observed(16). Notably, in this and other studies the addition of DNA polymerase theta inhibitors were shown to mitigate such kilobase-scale deletions(7,17). Moreover, AZD7648 has been shown to increase the occurrence of translocations, even when a single locus was targeted, highlighting the broad impact on genome architecture(14). These findings demonstrate that HDR enhancement via DNA-PK inhibition can be accompanied by complex structural variations that are not detectable by standard PCR amplification and short-read sequencing, underscoring the risk of large-scale genomic alterations(8).

While AZD7648 represents a powerful strategy to improve CRISPR-mediated gene correction, these studies highlight the critical need for comprehensive genomic assessment when applying DNA-PK inhibitors in both research and therapeutic contexts. Strand-seq is a single-cell DNA template strand sequencing technique that exploits the directionality of DNA strands inherited during replication to generate strand-specific read alignments for each chromosome in individual cells(18–21). Because each sister chromatid retains its parental template strand, inversions, deletions, duplications and translocations each produce characteristic shifts in read orientation that are detectable at single-cell resolution, without requiring long reads or paired samples. This strand-specific signal makes Strand-seq uniquely sensitive for detecting SVs that are missed by standard short-read sequencing, which lacks the directionality needed to distinguish rearrangements from mapping artefacts(18,20).

To investigate whether AZD7648-enhanced genome editing affects large-scale genome integrity in primary human HSPCs, we designed a multivariate strategy combining Strand-seq and long-read nanopore sequencing to assess both local ON-target and global structural variations in human HSPCs. This combination allowed precise mapping of chromosomal aberrations including inversions, deletions, duplications, and translocations as well as on-target large deletions at *DNMT3A* and *SRSF2*. Our results show that Cas9-mediated editing in general is sufficient to drive most large on-target deletions which, as observed by others(15,16), could be mitigated by PolQ inhibition. Importantly, the effect of AZD7648 was modest compared to editing in general, highlighting the need for thorough functional and genomic assessment of any potential edited cell therapeutic product.

## Results

### CRISPR editing includes large deletions at on-target loci which is increased by AZD7c48, but can be mitigated with PolQ inhibition

Initial work reporting large ON-target deletions used read length from nanopore sequencing as a proxy for large deletions(16). This risks the over-estimating large deletion frequency as truncated reads, would be counted as large deletions, a common issue in long-read sequences(19,22). As this strategy still makes use of PCR to amplify the targeted region, all complete sequences necessarily have at least the primer binding site on either end. It should thus be possible to unambiguously determine whether a given read was truncated due to technical factors or deletion. As such, we developed a new pipeline which anchors reads at both ends prior to determining large deletion abundance, thereby removing any truncated reads prior to quantification (Figure S1A, https://github.com/djhfknapp/Nanopore-Large-Deletion-Analysis). This bilateral anchoring approach provides a more stringent measure of large deletion frequency than methods relying on single-end read anchoring. Our pipeline was first validated against a synthetic dataset containing different large deletions of known size and frequency to confirm its accuracy for the application of interest (Figure S1B). When applied to samples with varying combinations of editing and DNA repair modulators on primary CD34+ HSPCs from human umbilical cord blood at the SRSF2 and DNMT3A loci, we were able to detect large deletion of varying sizes which centred on the Cas9 target site as expected (Figure 1A, S2, S3). Confirming the importance of bilateral anchoring, we observed a clear drop in coverage towards the end of the PCR product, even in unedited samples (Figure S1C, S4, S5). Beyond large deletion quantification, our pipeline also assesses on-target editing efficiency by reporting the percentage of reads carrying small indels or substitutions within reads that do not contain a large deletion. Importantly, these measures were highly correlated with those obtained by ICE analysis of the Sanger sequencing traces across all replicates and conditions (R² = 0.968; Figure S1D). This confirms that our pipeline provides an integrated readout including classical editing efficiency as well as large deletions outcomes.

**Figure 1.**
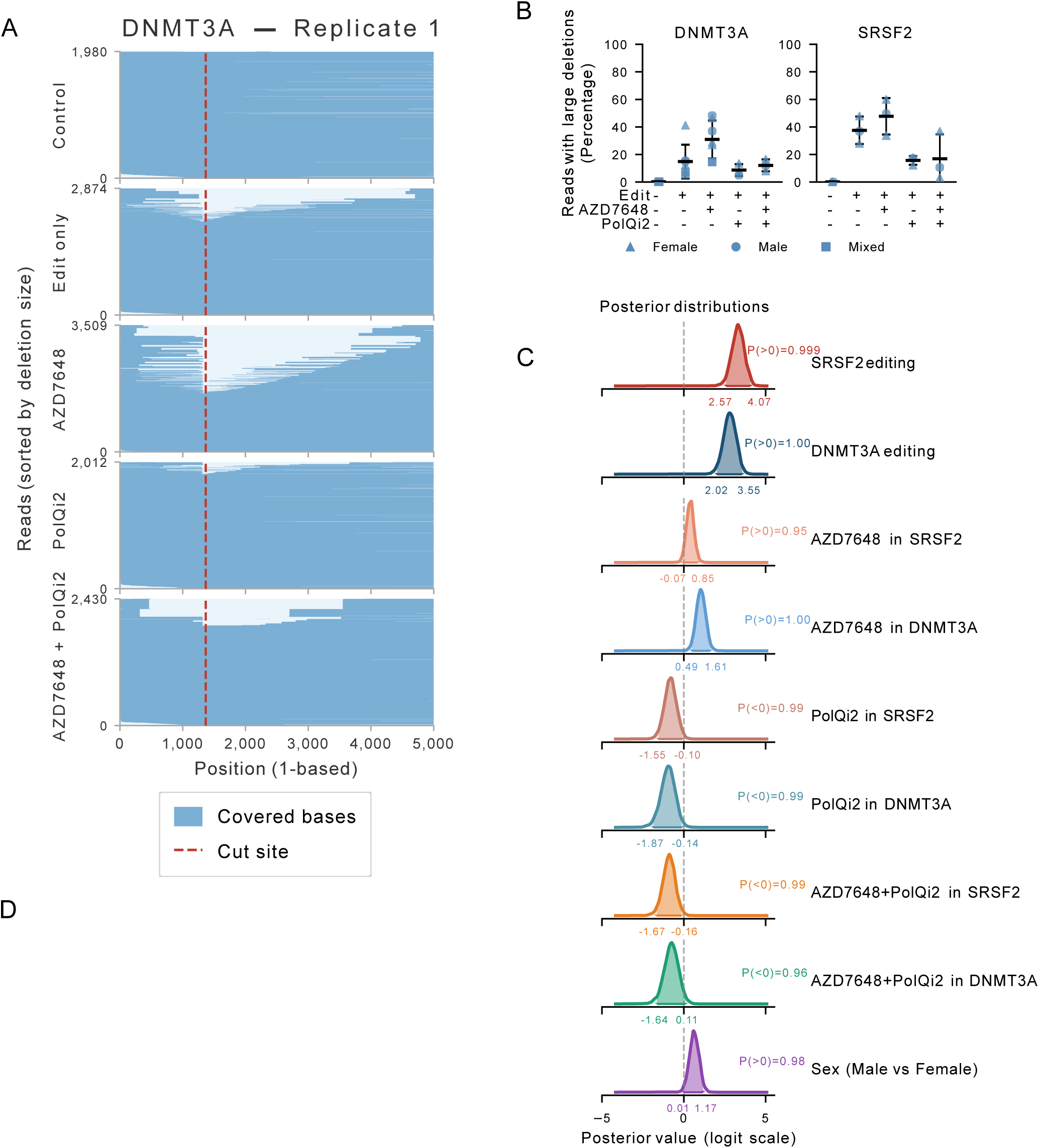
Large deletions at CRISPR-edited loci are suppressed by PolQ inhibition. Primary CD34+ hematopoietic stem and progenitor cells (HSPCs) from human umbilical cord blood were edited with Cas9 targeting the DNMT3A R882 or SRSF2 P95 locus under five conditions: unedited control, editing alone, editing with the DNA-PKc inhibitor AZD7648, editing with a PolQi2, or editing with both molecules. Individual male or female donor cords were used, with mixed-sex pools included in some replicates. On-target amplicons spanning each cut site were amplified by PCR (DNMT3A: 5,137pb; SRSF2: 4,188bp), barcoded, pooled, and subjected to Nanopore long-read sequencing. Reads were processed using our bilateral anchoring pipeline requiring read coverage on both sides of the cut site to ensure accurate large deletion quantification (see Supp Fig 1). **(A)** Representative deletion size distributions, sorted by large deletion size, from Nanopore long-read sequencing at the DNMT3A loci for replicate 1 detected across samples. Anchored reads are shown in blue, cas9 target site as a red dashed line, number of reads per condition is indicated on the top left Y axis. **(B)** Percentage of Nanopore sequencing reads carrying large deletions (>50 bp) for DNMT3A (5137 bp amplicon) and SRSF2 (4188 bp amplicon) across five experimental conditions: unedited control (condition Edit (-), AZD7648 (–)and PolQi2 (–) are considered as Control), editing alone (Edit (+)), editing with AZD7648 (AZD7648 (+)), editing with PolQ inhibitor (PolQi2 (+)), and editing with both AZD7648 and PolQi2 (AZD7648(+) and PolQi2 (+)). Individual data points represent biological replicates; horizontal bars indicate the mean; error bars represent ±1 SD. Marker shape corresponds to donor sex (triangle = female, circle = male, square = mixed). **(C)** Posterior probability density distributions for mixed-effect parameters from the Bayesian mixed-effects Beta regression model of large deletion frequency (proportion of reads with large deletions). The model was fitted with the formula: *prop* ∼ *(SRSF2 + DNMT3A) × AZD7648 × PolQi2 + Sex + (1 | Replicate)*, using a Beta likelihood with logit link function. SRSF2 and DNMT3A are binary fixed effects encoding gene-specific editing (reference: unedited control for each gene); AZD7648 and PolQi2 are binary fixed effects encoding drug treatment (reference: editing-only condition, no molecule); Sex is a fixed effect (reference: Female); and Replicate is a random intercept accounting for cord blood donor variability. Weakly informative priors were used: normal(0, 2) for fixed effects on the logit scale, Student-t(3, −4, 3) for the intercept, half-normal(0, 1) for random effect standard deviations, and gamma(0.01, 0.01) for the Beta precision parameter φ. Parameters shown represent composite contrasts of biological interest: editing effects (SRSF2 and DNMT3A vs. their respective unedited controls), total drug effects per locus (e.g. AZD7648 in SRSF2 = b₁AZD7648 + b₂SRSF2:AZD7648, vs. editing-only baseline), net combination effect (AZD7648+PolQi2 vs. editing-only), and the sex effect (Male vs. Female). All parameters are displayed on the log-odds (logit) scale; a dashed vertical reference line is shown at 0 (null effect). Positive values indicate an increase in large deletion frequency; negative values indicate a decrease. Shaded regions indicate the full posterior distribution (light) with the 95% highest density interval (HDI) highlighted (dark); HDI bounds are annotated numerically. Parameters whose 95% HDI excludes 0 indicate effects with high posterior certainty (posterior probability > 0.95). **(D)** Average size (bp) of large deletions per replicate across edited conditions for DNMT3A (5137 bp amplicon) and SRSF2 (4188 bp amplicon). Control condition excluded as large deletion sizes are not informative in unedited samples. Visual encoding as in (B).

Consistent with prior reports, editing drastically increased the proportion of reads carrying large deletions compared to unedited controls across both loci (Figure 1B) and this was not correlated with overall editing efficiency (R² = 0.021; Figure S6A). Bayesian mixed-effects modelling confirmed the association between editing and large deletion formation with high certainty (P(>0) = 1.00 at both loci; Figure 1B-C). AZD7648 treatment showed modest but consistent increase in large deletion frequency above the edited baseline at both loci (P(>0) = 0.95 for SRSF2 and P(>0) = 1.00 for DNMT3A; Figure 1B-C), consistent with recent reports that DNA-PKcs inhibition promotes alternative end-joining pathway activation(15,16). Similar results were also observed in an additional SRSF2 cohort measured over a shorter amplicon, though the overall frequency of large deletions detected was reduced in that cohort, consistent with missing the largest fraction of deletions which would have extended past the outside of even this longer than standard amplicon (Figure S6B-D). Notably, male cord donors showed a higher baseline rate of large deletions compared to female donors (Figure1B-C), suggesting a potential sex-dependent influence on DNA repair at Cas9 cut sites, though this would need to be confirmed with a larger dataset. Also consistent with previous reports(7,16), PolQ inhibition strongly mitigated large deletion rates, both when administered alone and in combination with AZD7648 (P(<0) = 0.99 for both locus; Figure 1B-C), confirming PolQ-mediated alternative end-joining as a major driver of large deletion formation at Cas9 cut sites, and reinforcing that PolQ inhibition represents a practical strategy for reducing on-target large deletion formation. As expected, AZD7648 substantially increased editing efficiency at both loci (∼85–90% vs. ∼47% for the p53 siRNA reference), while PolQi2 alone did not substantially alter editing efficiency above baseline, and its combination with AZD7648 did not improve efficiency beyond AZD7648 alone (Figure S1E), confirming that the modulation of large deletion frequency across conditions reflects repair pathway context rather than differences in overall editing rate.

We next examined whether these conditions also affect the size of large deletions among reads in which they occurred (Figure 1D). AZD7648 showed a modest positive trend in DNMT3A-edited cells (P(>0) = 0.75) but no directional trend in SRSF2-edited cells (P(>0) = 0.55)) (Figure S6E). PolQi2 showed a negative trend in both SRSF2 (P(<0) = 0.83)) and DNMT3A (P(<0) = 0.69)), suggesting smaller size deletions when PolQi2 is added (Figure S6E). Interestingly SRSF2 showed a slight trend towards smaller deletions compared to DNMT3A (P(<0) = 0.63), though in all cases the 95% highest density intervals (HDIs), the Bayesian analogue of a confidence interval, crossed zero (Figure S6E). Sex had no detectable influence on average deletion size (P(<0) = 0.54)), indicating that the observed effects of editing and small-molecule treatment are consistent across male and female donors (Figure S6E). Taken together, these results suggest that the experimental conditions primarily modulate the frequency of large deletion-bearing reads rather than the characteristic size of the deletions themselves.

To determine whether MMEJ contributes to CRISPR-induced deletions at these loci, we applied a read-confirmed microhomology classification pipeline (MH ≥ 5 bp) to all nanopore sequencing reads spanning the DNMT3A R882 and SRSF2 P95 cut sites across all conditions (Figure S6F). At the SRSF2 locus, only a single confirmed MMEJ event was detected across all samples and conditions, consistent with the near-absence of ≥ 5 bp microhomology positions in the SRSF2 amplicon reference sequence, precluding any quantitative assessment. At DNMT3A, MMEJ-associated deletions were detected at uniformly low frequencies across all conditions (Edit: 0.011 ± 0.007%; AZD7648: 0.008 ± 0.004%; PolQi2 and PolQi2+AZD7648: no confirmed events), with no condition-dependent trend observed (Figure S6F). These results indicate that MMEJ is not the main alternative to CRISPR-induced deletions at either locus under these experimental conditions, and that large deletion formation is instead driven predominantly by PolQ-dependent TMEJ, consistent with the reduced large deletion frequency observed with PolQ inhibition.

### Strand-seq of single cells from primary human cells in an HSC-enriched phenotype

Given the observed local effects, we next wanted to test whether AZD7648 and editing in general would give rise to structural variants across the rest of the genome. For this we made use of Strand-seq. CD34+ cells were nucleofected with SRSF2 and DNMT3A ssDNA donors in the presence or absence of AZD7648, and either in single editing or simultaneously at both loci in the presence of AZD7648. Following nucleofection, the HSC enriched population (CD34+CD45RA-CD90+) were isolated by flow sorting and grown for 7 to 9 days to ensure sufficient cells passing through S-phase to allow for Strand-seq (Figure 2A). All Strand-seq profiles thus represent the editing outcomes present in surviving HSC-enriched clones. We obtained high-quality profiles from 961 HSC-derived cells across six experimental conditions: Control, SRSF2 and DNMT3A editing alone without AZD7648, SRSF2 and DNMT3A editing alone with AZD7648 and dual editing of SRSF2 and DNMT3A simultaneously with AZD7648 (Figure 2A). The genome-wide strand-state profiles produced by Strand-seq allow, for each single cell, assessment of the Watson and Crick read distribution across all chromosomes, from which both the per-chromosome strand state and the donor sex can be computationally determined from Y chromosome coverage (information used in all downstream analyses) (Figure 2C-D). Finally, Strand-seq confirmed the expected per-chromosome strand states and enabled genome-wide SV detection at single-cell resolution such as SCE, deletion and duplication (Figure 2E).

**Figure 2.**
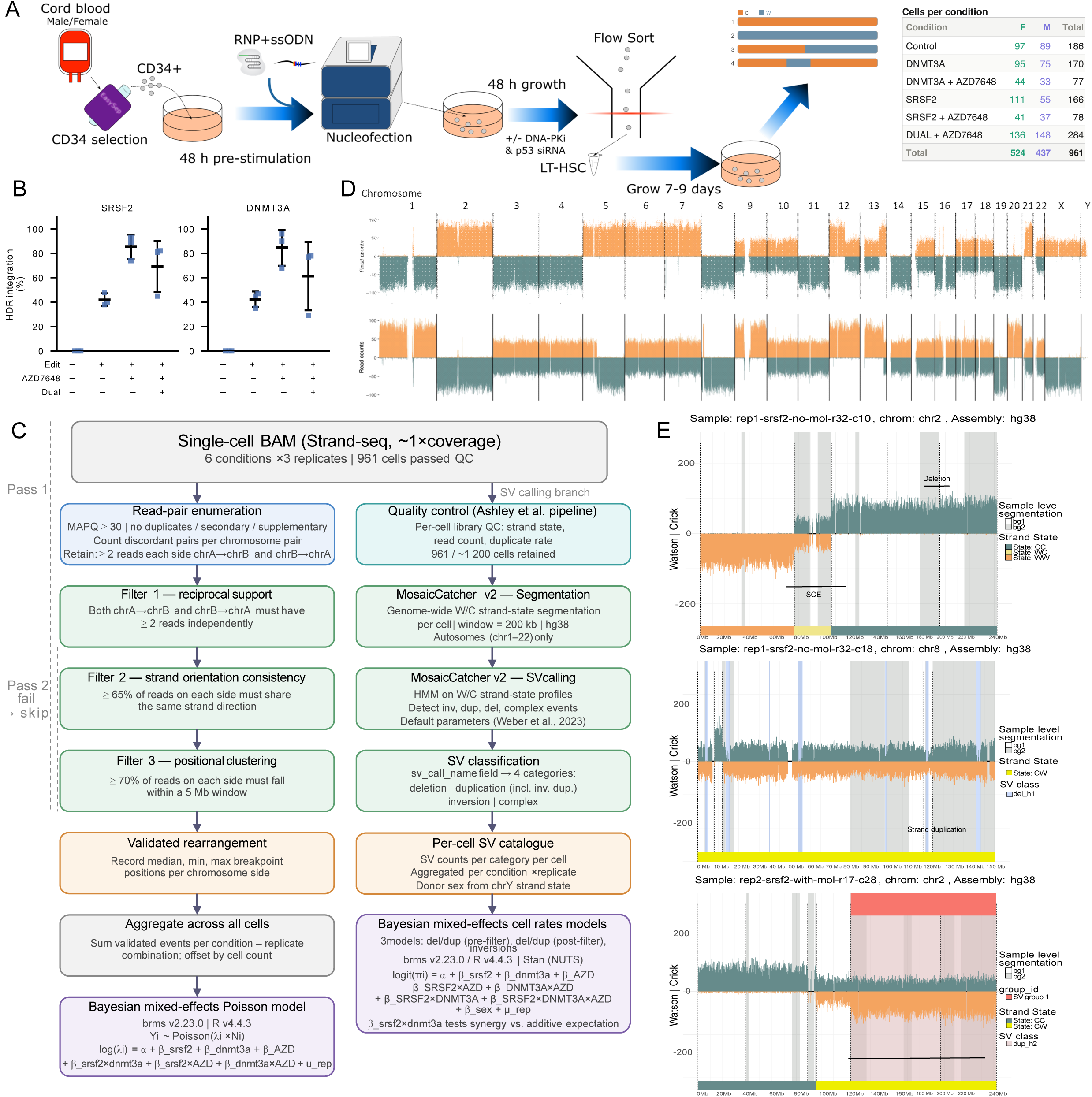
Experimental workflow and bioinformatic pipeline for single-cell Strand-seq analysis. **(A)** Schematic of the experimental workflow. CD34+ hematopoietic stem and progenitor cells were isolated from pulled cord blood cells (male and female donors) by CD34 selection using EasySep. Cells were pre-stimulated for 48 hours before nucleofection with RNP+ssODN complexes targeting SRSF2 and/or DNMT3A in the presence or absence of the molecule AZD7648. Following nucleofection, cells were cultured for 48 hours after which long-term hematopoietic stem cells (LT-HSCs) were isolated by flow sorting using the antibody panel CD34+CD45RA-CD90+CD49c+. Sorted cells were grown for 7–9 days to reach a sufficient number of cells before being sent for single-cell Strand-seq library preparation. **(B)** HDR integration efficiency (%) at the *SRSF2* and *DNMT3A* loci across the four experimental conditions used for Strand-seq analysis: unedited control (Ctrl), single-locus editing without AZD7648 (no mol), single-locus editing with AZD7648 (AZD), and simultaneous dual-locus editing with AZD7648 (Dual). Integration efficiency was measured by ICE analysis of Sanger sequencing traces, expressed as the percentage of reads carrying the intended HDR template sequence. Individual data points represent biological replicates (n = 3); horizontal bars indicate the mean; error bars represent ±1 SD. **(C)** Bioinformatic analysis pipeline. Single-cell BAM files were processed through two parallel branches. The left branch describes a custom translocation detection pipeline: read pairs were enumerated with quality filters (MAPQ ≥ 30, no duplicates/secondary/supplementary reads), retained if both orientations of a discordant chromosome pair had ≥ 2 supporting reads (Pass 1), and further filtered for strand orientation consistency (≥ 65% concordance) and positional clustering (≥ 70% of reads within a 5 Mb window, Pass 2). Validated rearrangements were aggregated per condition–replicate and modelled using a Bayesian mixed-effects Poisson model implemented in brms (v2.23.0, R v4.4.3), with binary predictors for SRSF2 editing, DNMT3A editing, and AZD7648 treatment, a cell-count offset, and a replicate random intercept. The right branch describes SV and SCE calling: cells passing quality control with the Ashley et al. pipeline (961/∼1,200 cells retained) were processed through MosaiCatcher v2 (Weber et al., 2023; hg38, 200 kb windows, autosomes chr1–22). SV calls were classified into four categories (deletion, duplication, inversion, complex) and modelled using five independent Bayesian mixed-effects count models (one per SV type plus total SV burden), incorporating editing status, AZD7648 treatment, donor sex, and a replicate random intercept. **(D)** Genome-wide Strand-seq read-count profile for a representative single cell, displayed across all autosomes and sex chromosomes. Orange and teal bars represent Crick and Watson strand read counts, respectively. The upper panel shows raw read counts; the lower panel shows the MosaiCatcher segmentation output. **(E)** Representative single-chromosome MosaiCatcher output plots for three cells illustrating different SV classes detected in the dataset. Each panel shows Watson (orange) and Crick (teal) read counts along the chromosome, with the strand-state segmentation overlaid. Coloured highlights indicate called SVs: a SCE (Strand WW, WC and CC, top panel, chr8), a hemizygous deletion (del_h1, middle panel, chr8), and a homozygous duplication spanning a large chromosomal region (right panel, chr2). Strand states (CC, WC, WW, CW) and SV group identifiers are indicated in the legend.

### Reciprocal translocations are rare but increased following CasG-editing

To assess whether individual or simultaneous edits in the presence or absence of AZD7648 affected reciprocal translocation occurrence we developed a custom genome-wide detection pipeline (Figure 2C, https://github.com/Fanny-Mei/scReciprocalTransloc). Existing SV callers applied to Strand-seq data are optimized for within-chromosome events and events that spread across different cells such as clones and do not directly report inter-chromosomal rearrangements from single-cell read data. Our custom pipeline instead leverages the strand-directionality of Strand-seq reads to detect reciprocal inter-chromosomal events at the single-cell level by requiring simultaneous reciprocal read evidence on both partner chromosomes, providing a complementary layer of genome-wide structural surveillance. By requiring simultaneous reciprocal read evidence, strand orientation consistency, and positional clustering, we identified 90 high-confidence inter-chromosomal rearrangement events across conditions. The circos plots reveal that cells which had been simultaneously edited at both the SRSF2 and DNMT3A loci displayed the highest number of high-confidence rearrangement events and the broadest genome-wide distribution of rearrangement partners, with one candidate event localizing within 5Mb of the SRSF2 cut site on chromosome (Figure 3A). Notably, 89 of the 90 translocation events did not involve the two targeted loci, suggesting that Cas9-induced breaks contribute to a genome-wide increase in translocations rather than exclusively site-specific ones.

**Figure 3.**
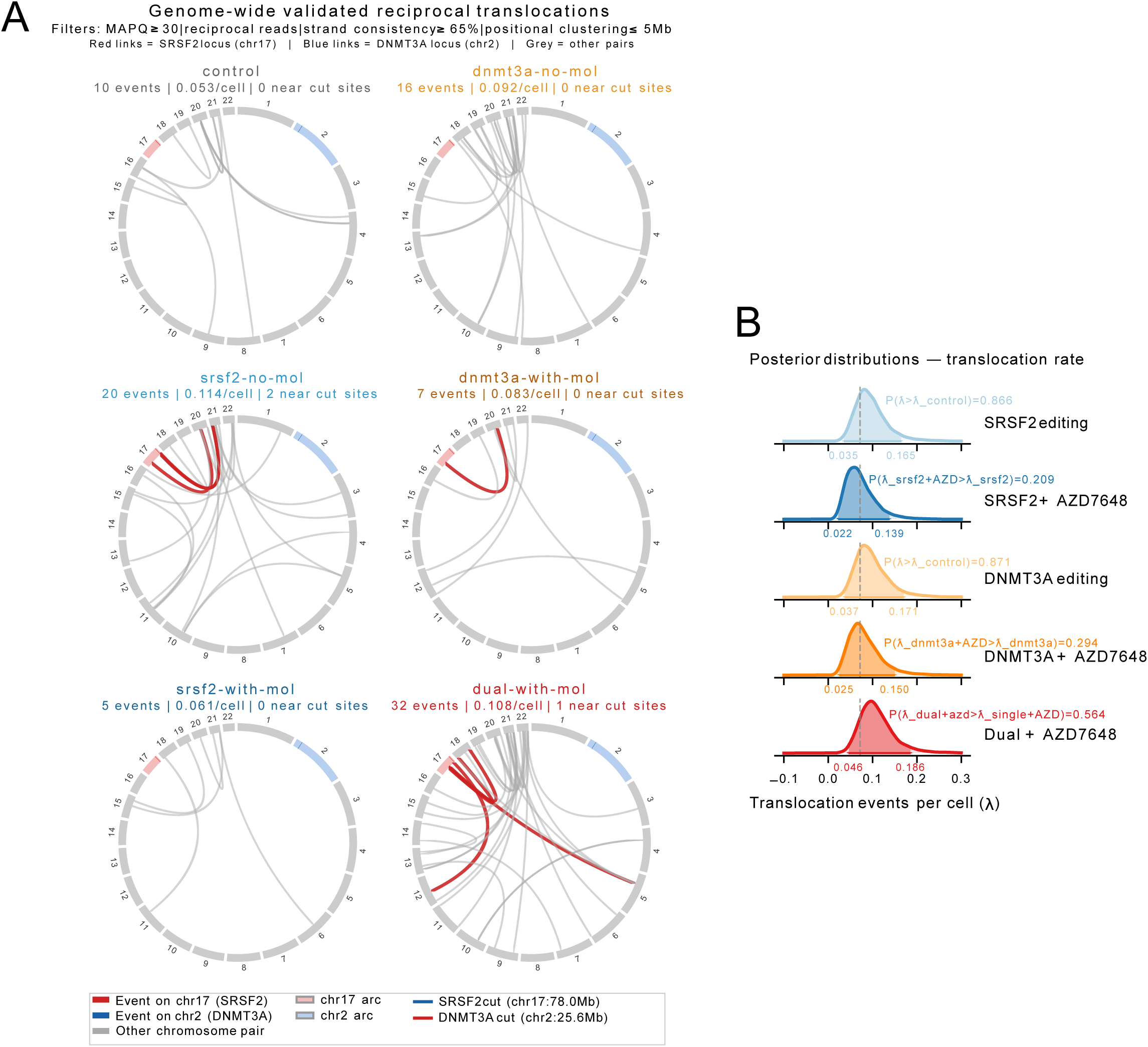
Genome-wide detection of reciprocal structural translocations in CRISPR-edited hematopoietic stem and progenitor cells. **(A)** Circos plots depicting genome-wide validated reciprocal structural rearrangements detected by pseudo-bulk Strand-seq analysis across six experimental conditions. Each chromosome is represented as an arc around the circle, with chromosome 17 (SRSF2 locus) highlighted in pink and chromosome 2 (DNMT3A locus) in light blue. Colored links connect chromosome pairs showing validated reciprocal discordant read pairs in the same cell. Red links indicate events where at least one breakpoint falls within 5 Mb of a CRISPR cut site. Red and blue rectangles on the chromosome arcs mark the SRSF2 P95H (chr17:78,015,826) and DNMT3A R882H (chr2:25,654,947) guide RNA cut sites, respectively. Events were validated by four simultaneous filters: (i) minimum mapping quality MAPQ ≥30, (ii) reciprocal read evidence on both chromosomes (≥2 reads per side), (iii) strand orientation consistency (≥65% of reads on each side sharing the same strand direction), and (iv) positional clustering of reads within a 5 Mb window. The total number of validated events, the normalized rate per cell, and the number of events near CRISPR cut sites are indicated above each circos plot. **(B)** Bayesian posterior distributions of reciprocal rearrangement rates (λ, events per cell) per condition, estimated from a Bayesian mixed-effects Poisson model (brms, R) with SRSF2 editing, DNMT3A editing, and AZD7648 as binary predictors with all pairwise interactions, and replicate as a random intercept. The model formula is: n_events ∼SRSF2*DNMT3A*AZD7648 + offset(log(n_cells)) + (1|replicate). Each panel shows the full posterior density (light fill), the 95% highest density interval (HDI, darker fill), and the posterior mean (vertical line). The dashed grey line indicates the control posterior mean as a reference. Posterior sampling used NUTS with 4 chains of 4 000 draws each (2 000 warmup, adapt_delta = 0.95). Weakly informative priors: intercept Normal(-2.3, 1.0) coefficients Normal (0, 0.5), random effect SD half-Normal (0.5).

To compare rearrangement rates across conditions while accounting for replicate variability and the factorial experimental design, we fit a Bayesian mixed-effects Poisson model, with SRSF2 editing, DNMT3A editing, and AZD7648 addition as binary predictors, all pairwise interactions, and replicate as a random effect (Figure 3B). In this framework, dual editing is not an independent condition but rather the interaction term between SRSF2 and DNMT3A editing, allowing us to directly test whether simultaneous cuts cause more rearrangements than the sum of individual effects. Editing at either locus alone was sufficient to elevate rearrangements rates above non-edited controls, with moderate evidence for both SRSF2 (P(λ > λ_control) = 0.866, and DNMT3A single edits (P(λ > λ_control) 0.871; Figure 3B). Simultaneous editing at both loci showed the strongest increase relative to control (P(λ > λ_control) = 0.933; Figure 3B). The SRSF2-DNMT3A interaction term was positive but did not reach strong evidence (P(λ_dual+AZD > λ_single+AZD) = 0.564), indicating that the elevated burden in dual-edited cells is consistent with an additive combination of the two individual editing effects. Surprisingly, the addition of AZD7648 did not increase reciprocal rearrangement rates, and in both the SRSF2 and DNMT3A interactions appeared to decrease the probabilities of these events (P(λ_srsf2+AZD > λ _srsf2) =0.209, P(λ_dnmt3a+AZD > λ _dnmt3a) =0.294). Taken together, these results suggest that CRISPR editing induces measurable but low rates of reciprocal translocations in HSPCs, that this is increased additively with additional edits, and that AZD7648 does not further aggravate but rather shifts the repair outcome away from reciprocal rearrangements and toward large deletions(23,24). Consistent with this, translocation rate did not correlate with HDR integration efficiency across conditions and replicates (Figure S7F), further supporting that the rearrangement burden is driven by the number of Cas9-induced breaks rather than the efficiency of template-directed repair.

### Genome-wide structural variant burden is amplified by simultaneous dual-locus CRISPR editing

To further assess genome-wide chromosomal integrity we next measured strand state, deletion, duplication and inversion events across conditions on our Strand-seq data (Figure 4A). Conditions included an unedited control (Ctrl), single-locus editing of *SRSF2* or *DNMT3A* without AZD7648, single-locus editing with AZD7648 (SRSF2 + AZD and DNMT3A + AZD), and simultaneous dual- locus editing of both *SRSF2* and *DNMT3A* with AZD7648 (Dual + AZD) (Figure 2A). To confirm successful editing across all conditions used for Strand-seq analysis, HDR integration efficiency was assessed by Sanger sequencing at both the SRSF2 and DNMT3A loci prior to cell sorting. As expected, AZD7648 substantially increased integration efficiency at both loci compared to editing alone (∼85–90% vs. ∼42%), and importantly, simultaneous dual editing of both loci with AZD7648 achieved comparable efficiencies at each locus, confirming that co-editing of SRSF2 and DNMT3A in the same cells is feasible under these conditions (Figure 2B).

**Figure 4.**
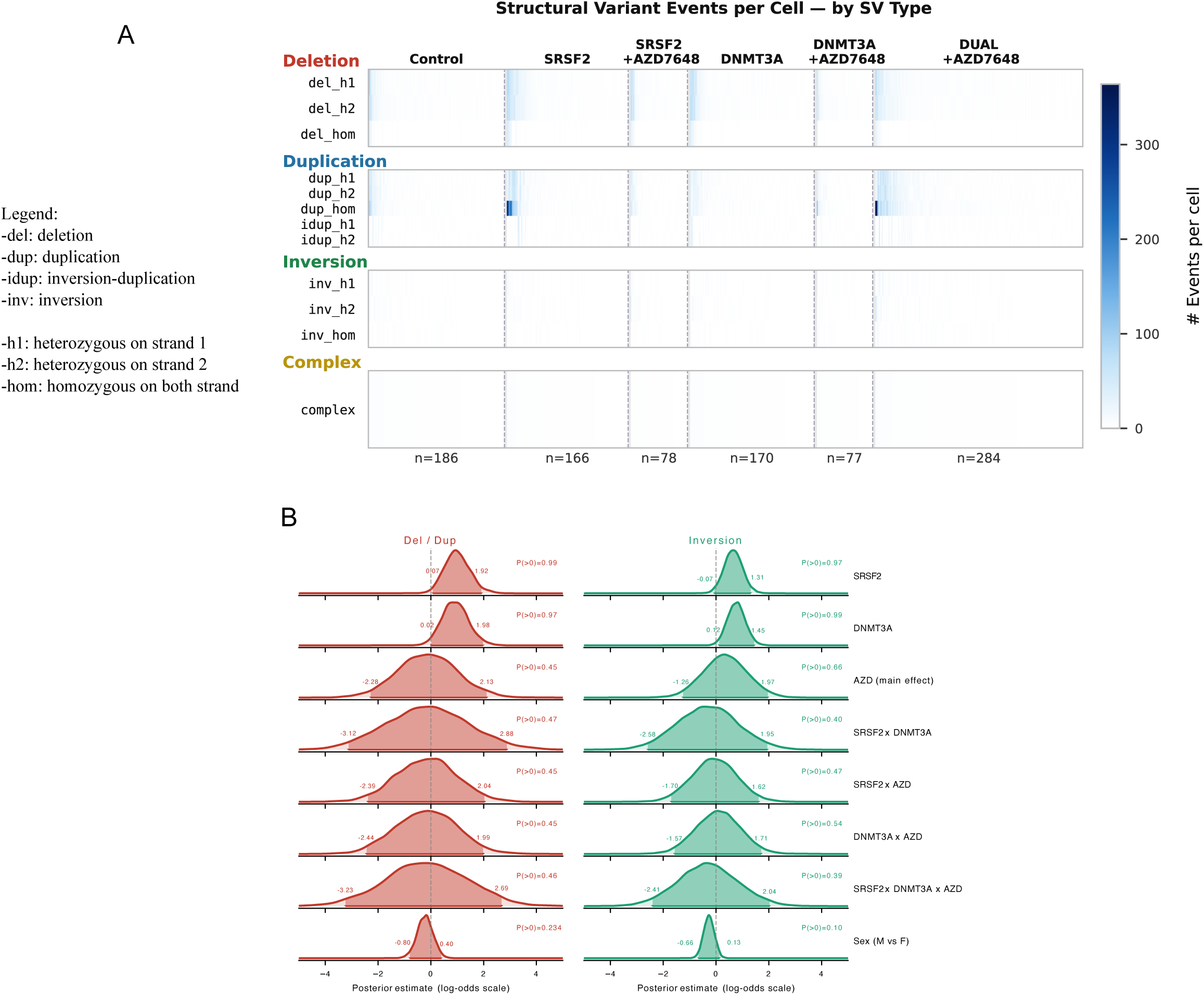
Genome-wide structural variant burden is amplified by dual-locus CRISPR editing. **(A)** Heatmap of per-cell SV event counts across all conditions and SV subtypes. Each column represents one single cell, sorted by total SV count within each condition block (separated by dashed vertical lines). Colour intensity reflects the number of events per cell on a linear scale (white = 0, dark navy = maximum). SV subtypes are grouped by class along the y-axis: deletions (del_h1, del_h2, del_hom), duplications (dup_h1, dup_h2, dup_hom, idup_h1, idup_h2), inversions (inv_h1, inv_h2, inv_hom), and complex events. n = 961 cells; 34,787 SV events; 3 independent replicates. h1 stands for strand 1, h2 for strand 2 and hom for homozygous on both strands. **(B)** Bayesian mixed-effects analysis of del/dup and inversion cell-positive rates. Posterior density plots showing the effect of each experimental contrast on the log-odds of a cell carrying at least one event, for del/dup (left) and inversion (right) models separately. The shaded area represents the full posterior density; the dashed vertical line marks log-odds = 0 (no effect). The horizontal line below each density indicates the 95% highest density interval (HDI); the filled circle marks the posterior median. Annotated values show the posterior median and posterior probability P(FC>1) or P(FC<1) as appropriate for the direction of the contrast. The del/dup model was fitted on events spanning ≥10% of the chromosome arm length, with an arm-weakness filter applied to exclude recurrently called control arms, and downstream removal of control-like and chromosome-spread cells. The inversion model was fitted on events filtered using the arm-weakness approach with a shared-control blacklist at reciprocal overlap ≥0.6. Both models use a Bernoulli (binomial) likelihood with a logit link, with random intercepts per biological replicate, fitted using brms (R interface to Stan).

To quantify the effect of editing genotype, AZD7648 treatment, sex, and their interactions on SV burden, we fit Bayesian mixed-effects models under two filtering strategies for deletion and duplication events. These two complementary size-filtering strategies were applied to deletion and duplication calls to assess result robustness and to distinguish genuine somatic events from technical noise. A lenient 5 Mb threshold was used as an initial sensitivity analysis, as it captures the broadest range of potential events while remaining above the resolution limit of the MosaiCatcher 200 kb binning. However, this threshold retains a higher proportion of small, recurrent events that appear in control cells and likely reflect systematic mapping artefacts at repetitive or low-complexity regions. A more stringent filter, requiring events to span at least 10% of the chromosome arm length combined with an arm-weakness filter excluding chromosome arms recurrently called in control cells, was therefore applied as the primary analysis to reduce false positives while preserving large, condition-enriched events. For inversions, the arm-weakness filter alone was applied with no size restriction, as inversions naturally occur in cells (25,26). Regardless of the filtering strategy applied, both models showed clear effects of SRSF2 and DNMT3A editing on del/dup burden (lenient filter: P(FC>1) = 1.00 for both; stringent filter: P(FC>1) = 0.99 and P(FC>1) = 0.97, respectively), with no strong evidence for an effect of AZD7648 (P(FC>1) = 0.38 and P(FC>1) = 0.45, respectively) or synergistic effects of dual editing or AZD7648 combination, though additive effects of individual mutations were present. No clear sex effect was observed (P(FC>1) = 0.234) (Figure 4B and Figure S7A). A total of 184 inversion-positive cells were observed across conditions, and inversions showed similar trends with clear editing effects (SRSF2: P(FC>1) = 0.97; DNMT3A: P(FC>1) = 0.99), though in this case there was evidence for a modest increase related to AZD7648 exposure (P(FC>1) = 0.66, FC = 1.41) (Figure 4B).

Taken together, these results indicate that editing at SRSF2 or DNMT3A broadly elevates both large-scale del/dup and inversion burden, while AZD7648 reinforces only the inversion events, pointing to a context-dependent shift in DNA repair pathway usage upon NHEJ inhibition. Deletion and duplication burden can be attributed more to a general Cas9 off-target effect than to AZD7648 specifically, and neither del/dup nor inversion rates correlated with HDR integration efficiency across conditions and replicates (Figure S7G–H), indicating that SV formation is independent of the efficiency of on-target repair.

Sister chromatid exchange (SCE) frequency was also assessed to determine whether prior editing could alter long-term replication-associated DNA damage and repair activity (Figure S7B). SCE counts per cell were broadly comparable across all six conditions, with median values ranging from approximately 3 to 5 SCEs per cell. No condition showed a dramatic departure from the control distribution, and AZD7648 treatment did not substantially alter SCE frequency in either SRSF2- or DNMT3A-edited cells. This indicates that previous NHEJ inhibition by AZD7648 does not induce any lasting effects on replication fidelity. The stability of SCE counts across conditions further supports the interpretation that the structural variant landscape is shaped by transient, locus-specific repair events rather than a lasting genome-wide replication stress phenotype.

To determine whether SV events are randomly distributed across the genome or preferentially enriched at specific chromatin compartments, we next mapped all SV events to two genomic region classes defined from hg38 cytobands: euchromatin and heterochromatin (Figure S7C). Each SV event was assigned to a chromatin class based on maximum base-pair overlap with hg38 cytoband annotations collapsed into the two categories. For events spanning multiple cytobands, the class with the greatest cumulative overlap was used for assignment. Across all conditions, 96 to 100% of events occurred in euchromatic regions, with heterochromatin accounting for at most 3.8% of events in any condition. This euchromatic enrichment was observed in both edited and non-edited cells. In control cells, the vast majority of detected events are inversions, consistent with known polymorphic inversion variants that arise preferentially in open chromatin regions (17,18); deletion and duplication events in control cells are rare and largely filtered out by the arm-weakness filter, which was designed precisely to exclude such recurrent control-cell artefacts. In edited cells, the additional events include both inversions and del/dup events, with the latter showing the strongest condition-dependent increases above baseline. It should be noted that because retained SV events span large genomic segments (≥10% of chromosome arm length), individual events may visually overlap with heterochromatic cytobands in the ideogram (Figure 5A, Figure S8); however, chromatin class assignment was based on maximum base-pair overlap, and the majority of each event’s span falls within euchromatic regions. In edited cells, the additional DSB-induced events also predominantly affected euchromatic regions, consistent with the preferential accessibility of euchromatin to Cas9 and the higher density of replication-associated fragile sites in open chromatin compartments. Together, these results suggest that editing amplifies an existing chromatin-linked vulnerability rather than introducing a fundamentally distinct pattern of genome instability.

**Figure 5.**
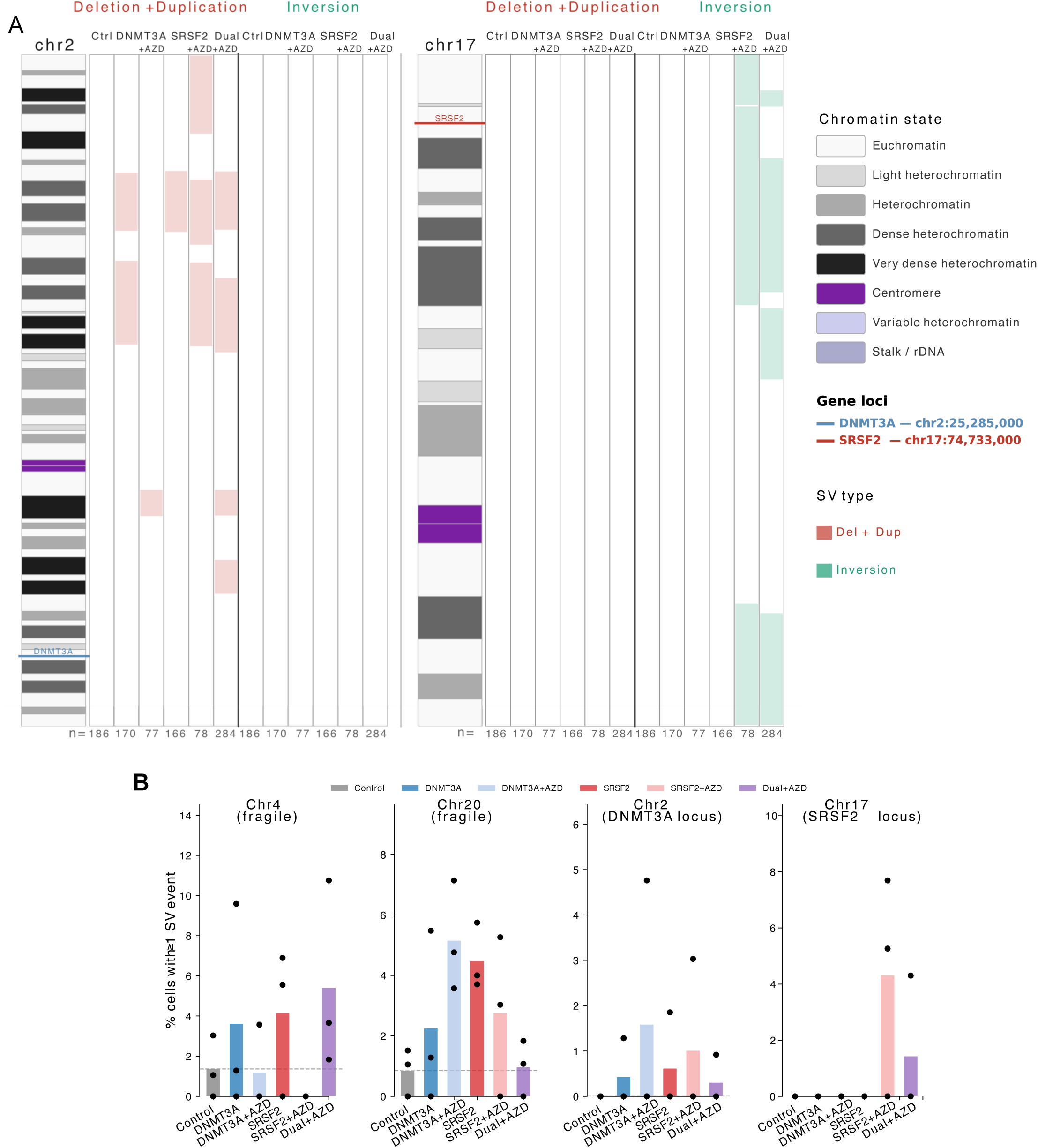
SV events are not primarily located at on-target sites but rather sparse across the genome. **(A)** Chromosome ideograms for chr2 (DNMT3A) and chr17 (SRSF2) are shown for each experimental condition (n = number of single cells pooled across three biological replicates), with all six conditions displayed side by side for each chromosome to facilitate direct comparison. Cytogenetic banding has been defined using: UCSC Genome Browser. Coloured horizontal lines indicate the genomic position of DNMT3A (chr2:25,285,000, blue) and SRSF2 (chr17:74,733,000, red). For each condition, two SV tracks are displayed to the right of the shared ideogram: deletions and duplications combined (red) and inversions (teal), separated by a vertical dark line. Each bar represents the full genomic span of an individual SV event. No overlapping events have been identified on these two chromosome, thus colour opacity gradient is not relevant in this panel. Only large structural variants representing at least 10% of the corresponding chromosome arm length were retained, and events overlapping centromeric and pericentromeric regions were excluded prior to visualization. SV calls were generated with the MosaiCatcher pipeline (hg38). **(B)** The percentage of cells carrying at least one validated SV event (del/dup or inversion) on chromosomes of interest across experimental conditions. Each panel shows one chromosome: Chr4 and Chr20 are known constitutively fragile chromosomes; Chr2 and Chr17 harbour the DNMT3A (2p23.3) and SRSF2 (17q25.1) editing loci respectively. Bar height indicates the mean percentage of cells affected across three biological replicates; each dot represents one replicate. The dashed grey line marks the control mean as a reference. No statistical testing was applied given the limited number of events per condition.

Mapping of large structural variant events across the genome revealed a non-uniform chromosomal distribution, with recurrent hotspots identified on several autosomes across conditions (Figure 5A, Figure S8). To assess whether Cas9 editing preferentially destabilizes known chromosomally fragile regions, we examined the per-cell SV burden on chromosomes 4 and 20, two chromosomes with well-documented structural instability (27–31), across all conditions using the filtered retained event set (Figure 5B). In control cells, chr4 and chr20 were affected in 1.6% and 1.1% of cells respectively, consistent with baseline fragility. Across edited conditions, chr4 event rates were elevated in DNMT3A (4.7%), SRSF2 (5.4%), and Dual+AZD edited cells (5.3%), representing a 3- to 4-fold increase over control (Figure 5B). Similarly, chr20 showed elevated rates in DNMT3A (2.9–5.2%) and SRSF2-edited cells (4.8%) compared to control (Figure 5B). These differences did not reach statistical significance by Fisher’s exact test, likely due to the limited number of events per condition, but the consistent directional enrichment across independent edited conditions suggests that Cas9-induced double-strand breaks may exacerbate the intrinsic fragility of these chromosomes. Notably, chr17 and chr2, the chromosomes harbouring the SRSF2 and DNMT3A editing loci respectively, showed condition-specific signals, with chr17 events detected exclusively in SRSF2+AZD (4.23 per 100 cells) and Dual+AZD cells (1.41 per 100 cells), and chr2 events modestly elevated in DNMT3A-edited conditions, potentially reflecting on-target large deletion or rearrangement events at the editing locus (Figure 5A-B).

To characterize the size spectrum of deletion events genome-wide, we examined deletion/duplication and inversion size distributions per condition across all retained events after filtering (Figure S7D). No significant differences in event size were detected between conditions for either deletion/duplication or inversions (Mann-Whitney U test), indicating that editing with or without the molecule AZD7648 do not systematically shift the size spectrum of validated events. The size distributions were broadly comparable across edited and non-edited cells, with deletion/duplication events spanning from ∼10 kb to >100,000 kb and inversions ranging from ∼10 kb to ∼10,000 kb across all conditions.

## Discussion

This study provides a comprehensive assessment of genome integrity following CRISPR-Cas9-mediated editing in human HSPCs, combining long-read nanopore sequencing for on-target large deletion quantification with Strand-seq for genome-wide structural variant detection. Our findings converge on several important observations with relevance to both the mechanistic understanding of CRISPR-induced DNA repair and the practical development of therapeutic genome editing strategies.

A central finding of this work is that large on-target deletions are predominantly a consequence of CRISPR-Cas9 editing itself, rather than a specific effect of DNA-PK inhibition with AZD7648. While AZD7648 did produce a modest but consistent increase in large deletion frequency above the edited baseline at both the SRSF2 and DNMT3A loci, this effect was substantially smaller than the increase attributable to editing alone. This is consistent with and extends recent reports demonstrating that large deletions are a general feature of Cas9-induced double-strand break repair and that AZD7648 does have an additive effect(7,16), and underscores the importance of assessing these outcomes regardless of whether pharmacological repair modulation is employed in genome editing strategy. The bilateral anchoring strategy used in our nanopore pipeline ensured that truncated reads, which could otherwise inflate deletion estimates, were excluded, providing a more accurate frequency measure than approaches relying on single-end anchoring.

Consistent with previous reports, PolQ inhibition substantially reduced on-target large deletion frequency without compromising overall editing efficiency, both when used alone and in combination with AZD7648. The absence of a dominant MMEJ signature at either locus, with very low frequencies of reads bearing canonical microhomology (≥5 bp) at deletion junctions, suggests that PolQ-dependent theta-mediated end joining (TMEJ), rather than classical MMEJ, is the primary alternative pathway responsible for large deletion formation at Cas9 cut sites under these conditions. This mechanistic distinction is important, as it clarifies why PolQ inhibition is effective and implies that the deletions rescued by this inhibitor are largely the product of a specific polymerase-dependent repair mechanism rather than a broad dysregulation of end joining.

At the overall genome level, our Strand-seq analysis revealed that structural variants are present at low but detectable frequencies in all edited conditions, and that the overall burden is induced by editing generally rather than specifically due to AZD7648 addition. The genome-wide SV landscape was shaped predominantly by SRSF2 editing, which drove substantial increases in deletion and duplication burden, while DNMT3A editing alone had a more modest effect. The reasons for this locus-specific difference are not immediately clear but may reflect differences in local chromatin architecture or the genomic context surrounding each cut site. Importantly, the increase in deletion and duplication burden was attributable to Cas9 editing in general, with AZD7648 showing little evidence of an additional effect on large scale SVs.

Reciprocal translocations were detected at low frequencies across all conditions, confirming that Cas9-induced double-strand breaks can generate inter-chromosomal rearrangements in primary HSPCs, consistent with previous reports in multiplex editing strategies (10,11). Indeed, we observed that such translocations were greatly increased with multiple editing, suggesting increased numbers of double stranded breaks increase the recombination probability. In contrast to what was reported by Cullot et al., AZD7648 did not increase reciprocal translocation rates in our dataset; if anything, there was a trend toward reduced translocation frequency in AZD7648-treated cells. The discrepancy with Cullot et al. may reflect methodological and biological differences. CAST-seq, used in their study, preferentially detects translocations linked to the edited locus in bulk with high sensitivity, whereas Strand-seq used in ours captures genome-wide chromosomal rearrangements at the single-cell level. Consequently, the two approaches interrogate distinct aspects of genome instability and may differ in their sensitivity to specific classes of translocation events. In addition, different target loci were examined in the two studies, and both datasets suggest that locus-specific genomic context can substantially influence the spectrum of editing outcomes, including translocation formation.

Several limitations of this study should be considered. The detection of SVs at the genome-wide level by Strand-seq is dependent on the resolution afforded by both overall genome coverage and the 200 kb window size used for event calling. As such, sensitivity will be somewhat variable across cells, and smaller rearrangements may not be captured particularly in cells with lower coverage. Additionally, analyses were performed on cells expanded for 7–9 days after editing to enable sufficient numbers for Strand-seq, meaning that cells bearing highly deleterious rearrangements would have been selectively depleted from the population prior to analysis. This could potentially lead to an underestimate of the initial frequency of such events that could have been detected in other reports(15,16), though ultimately it is those cells that survive which would be potentially dangerous in a therapeutic setting which we would capture in our approach. Mechanistically, the trend we observe is coherent with the known biology of NHEJ inhibition: by suppressing the primary pathway that mediates end-joining between heterologous chromosome ends, AZD7648 may paradoxically reduce the probability of reciprocal translocations detectable at the chromosomal scale, while redirecting repair toward large on-target deletions. This interpretation is consistent with the redistribution of repair outcomes we observe across SV types, rather than a simple additive increase in unstable events. Additional editing events increased rearrangement rates in an approximately additive manner, consistent with the expectation that each additional double-strand break independently increases the probability of aberrant inter-chromosomal end joining.

From a translational perspective, these findings collectively suggest that the risk associated with HDR-enhanced CRISPR editing in HSPCs is present with any cut-based editing approach, and that the incremental risk attributable to AZD7648 specifically is relatively modest. The addition of PolQ inhibition represents a practical and effective mitigation strategy for on-target large deletions, consistent with its emerging role in precision editing protocols. None the less, the frequency of these events even under optimal scenarios means that some proportion of hemizygous cells are likely in any edited product and this should be carefully considered and large deletions assessed when using these techniques. Additionally, while still a minority, some cells bearing an abundance of structural abnormalities still arose and were viable at least for a week. This combined with the fact that most events were unique to only a single cell in the dataset means that they would likely be missed by most sequencing-based approaches. Functional assessments of the therapeutic cell population for clonal outgrowth may provide a powerful avenue to detect extremely rare but dangerous clones. Overall, our results reinforce the need to comprehensively assess any potential therapeutic cell product for potentially deleterious minor clones that may be introduced in the editing process.

## Supporting information

Key resources table

Significance Tests

Raw Data

## Acknowledgements

Funds for this work were provided by the IRIC Philanthropic funds from the Marcelle and Jean Coutu foundation and the Terry Fox Research Institute (Terry Fox New Investigator Award #TFRI 1118 and the Terry Fox New Frontiers Program Project Grant in Strategies to Divert Malignant Potentials in Acute Leukemia Award # 1135-03). FMCU was supported by a Cole Foundation Doctoral Award, a bourse d’excellence du programme de biologie moléculaire from the Université de Montréal and a PhD scholarship from the Institut de Recherche en Immunologie et en Cancérologie. The authors would like to thank Thomas Sontag and Michel Duval from the Banque de Recherche de Sang de Cordon at CHU Sainte-Justine and HemaQuebec for assistance with cord blood acquisition, the generous individuals who donated their cords for this project, Annie Gosselin and Angélique Bellemare-Pelletier from the IRIC Flow Cytometry facility for technical assistance with sorts, Raphaelle Lambert from the IRIC Genomics facility for technical assistance with Sanger sequencing. We thank Daniel Chan, Tiffany Leung, Yanni Wang, Mianne Lee and Trang Nguyen of the Strand-seq core facility at the BC Cancer Research Institute in Vancouver, BC for processing and sorting cultured hematopoietic cells and generating high quality Strand-seq data.

## CRediT Statement

- Cloarec-Ung FM: Conceptualization, Data curation, Formal Analysis, Investigation, Methodology, Project administration, Software, Validation, Visualization, Writing – Original draft, Writing – review and editing
- Sauvageau G: Resources, Writing – review and editing
- Sheppard HM: Investigation, Supervision, Writing – review and editing
- Lansdorp PM: Investigation, Resources, Software, Writing – review and editing
- Knapp DJHF: Conceptualization, Formal Analysis, Funding acquisition, Investigation, Methodology, Project administration, Software, Supervision, Visualization, Writing – Original draft, Writing – review and editing

## Data availability statement

Sequencing data is available through EGA with accession numbers EGAS50000002119, EGAS50000002120, and EGAS50000002121.

## Supplementary Figure Legends

**Supplementary Figure 1. Pipeline validation for large deletion detection at on-target site. (A)** Schematic overview of the Nanopore long-read sequencing pipeline for large deletion detection. Reads are aligned to the reference amplicon and required to anchor on both sides of the targeted cut site (bilateral anchoring). Reads that do not span the full amplicon are discarded prior to quantification. Reads with an aligned deletion exceeding 50 bp relative to the reference are classified as large-deletion reads. The pipeline simultaneously reports on-target editing efficiency as the percentage of reads carrying any indel at the cut site. **(B)** Validation of pipeline sensitivity across deletion sizes using a synthetic dataset. In silico-generated deletion alleles spanning the range of detectable deletion sizes were introduced at defined frequencies and processed through the pipeline, confirming accurate detection and quantification without systematic size-dependent bias. **(C)** Coverage profile across the PCR amplicon at a representative locus, llustrating the drop in read coverage toward the 3’ end characteristic of truncated Nanopore reads. These truncated reads, which cannot be assigned a true deletion size, are removed prior to large deletion quantification by the bilateral anchoring filter, ensuring that coverage drop-off artifacts do not inflate large deletion estimates. **(D)** Correlation between editing efficiency measured by Sanger sequencing (ICE analysis) and by the Nanopore pipeline across all replicates and conditions (R² = 0.968), validating the Nanopore pipeline as a reliable substitute for Sanger-based editing efficiency measurement. Each data point represents one replicate × condition × locus measurement; the solid line shows the ordinary least squares (OLS) regression fit; the shaded band represents the 95% confidence interval; the dashed line indicates the identity line (slope = 1). **(E)** Sanger sequencing-based editing efficiency (%) across all conditions at the SRSF2 and DNMT3A loci. Each data point represents one biological replicate; horizontal bars indicate the mean; error bars represent ±1 SD. Marker shape corresponds to donor sex (triangle = female, circle = male, square = mixed). AZD7648 substantially increased editing efficiency at both loci (∼85–90% vs. ∼47% for the p53 siRNA reference), while PolQi2 alone or in combination with AZD7648 did not substantially alter editing efficiency relative to AZD7648 alone.

**Supplementary Figure 2. Per read coverage panels for each replicate of batch 1 for large deletion assessment.** Representative deletion size distributions, sorted by large deletion size, from Nanopore long-read sequencing at the DNMT3A loci for replicate 2 and 3 and at the SRSF2 for replicate 1, 2 and 3 detected across samples. Only reads anchored on both sides of the cut site (bilateral anchoring) are included in the analysis. Anchored reads are shown in blue, cas9 target site as a red dashed line, number of reads per condition is indicated on the top left Y axis.

**Supplementary Figure 3. Per read coverage panels for each replicate of batch 2 for large deletion assessment.** Representative deletion size distributions, sorted by large deletion size, from Nanopore long-read sequencing at the DNMT3A loci for replicate 1, 2, 3 and 4 (Replicates 5, 6 and 7 have been discarded because of having less than a 1000 reads sequenced for the Control condition) and at the SRSF2 for replicate 1, 2, 3, 4, 6 and 7 (Replicate 5 has been discarded because of having less than a 1000 reads sequenced for the Control condition) detected across samples. Only reads anchored on both sides of the cut site (bilateral anchoring) are included in the analysis. Anchored reads are shown in blue, cas9 target site as a red dashed line, number of reads per condition is indicated on the top left Y axis.

**Supplementary Figure 4. Coverage profile for all replicates from batch 1.** Coverage profile across the PCR amplicon at a representative locus, illustrating the drop in read coverage toward the 3’ end characteristic of truncated Nanopore reads for both locus SRSF2 and DMT3A. These truncated reads, which cannot be assigned a true deletion size, are removed prior to large deletion quantification by the bilateral anchoring filter, ensuring that coverage drop-off artifacts do not inflate large deletion estimates.

**Supplementary Figure 5. Coverage profile for all replicates from batch 2.** Coverage profile across the PCR amplicon at a representative locus, illustrating the drop in read coverage toward the 3’ end characteristic of truncated Nanopore reads for both locus SRSF2 and DMT3A. These truncated reads, which cannot be assigned a true deletion size, are removed prior to large deletion quantification by the bilateral anchoring filter, ensuring that coverage artifacts do not inflate large deletion estimates.

**Supplementary Figure 6. Large deletion characteristics at CRISPR-edited loci: amplicon size effects, repair pathway contributions, and editing efficiency independence. (A)** Correlation between Nanopore-derived editing efficiency and the percentage of reads carrying large deletions across edited conditions (Ctrl excluded). Each data point represents one replicate × condition × locus measurement. The solid line shows the OLS regression fit; the shaded band represents the 95% confidence interval. The absence of correlation (R² = 0.021, p = 0.353) indicates that large deletion frequency is not a byproduct of editing rate but is determined by the specific repair pathway context imposed by each experimental condition. **(B)** Comparison of raw large-deletion frequencies detected with two SRSF2 PCR amplicons of different sizes (1157 bp and 4188 bp) across Ctrl, no molecule, and AZD conditions. Within each condition, the two amplicon sizes are plotted side by side. Colour encodes amplicon size (teal = 1157 bp; blue = 4188 bp); marker shape encodes donor sex (triangle = female, circle = male, square = mixed). The 4188 bp amplicon consistently yields higher detected large deletion frequencies, reflecting the greater sequence space available for deletion detection in longer amplicons. The two amplicons should not be compared quantitatively across batches. **(C-D)** Average size (bp) of detected large deletions for the two SRSF2 amplicons (1157 bp and 4188 bp) under no molecule and AZD conditions. The longer amplicon captures larger average deletion sizes, consistent with the interpretation that the 1157 bp amplicon truncates the detectable deletion size range and systematically underestimates both the frequency and magnitude of large deletion events. These data support the use of amplicons of at least ∼4 kb for accurate large deletion quantification at CRISPR-edited loci. (**E**) Posterior probability density distributions for mixed-effect parameters from the Bayesian mixed-effects Gaussian linear model of average large deletion size (bp). The model formula was identical to (Fig1C): *avg_del_size* ∼ *(SRSF2 + DNMT3A) × AZD7c48 × PolQi2 + Sex + (1 | Replicate)*, with an identity link function and a Gaussian likelihood. Fixed and random effects specifications and reference levels were identical to the frequency model. Weakly informative priors were specified on the base pair scale: normal(0, 500) for fixed effects, Student-t(3, 1500, 500) for the intercept (centred near the observed mean of 1,482 bp), half-Student-t(3, 0, 500) for the residual standard deviation, and half-normal(0, 300) for random effect standard deviations. Parameters shown are composite contrasts identical in structure to (Fig. 1C), displayed directly in base pairs (no transformation); a dashed vertical reference line is shown at 0 bp (null effect). Positive values indicate larger deletions; negative values indicate smaller deletions. Shaded regions and HDI annotations as in (Fig. 1C). No fixed-effect parameter reached a posterior probability > 0.95, indicating that experimental conditions modulate the frequency rather than the characteristic size of large deletions. **(F)** Frequency of CRISPR-induced deletion events bearing read-confirmed microhomology ≥ 5 bp at the DNMT3A R882 and SRSF2 P95 cut site junctions, expressed as percentage of total reads. Analysis was performed on all Nanopore reads across two independent experiments (no molecule and AZD7648: n = 9 replicates; PolQi2 and PolQi2+AZD7648: n = 3 replicates). Each dot represents one biological replicate; marker shape indicates donor sex (triangle = female, circle = male, square = mixed); horizontal bar = mean; error bars = ±1 SD.

**Supplementary Figure 7. AZD7648 does not substantially alter genome-wide structural variant burden, sister chromatid exchange frequency, or SV size distributions in CRISPR-edited HSPCs. (A)** Bayesian mixed-effects analysis of del/dup cell-positive rates under the initial 5 Mb size threshold (pre-filtering sensitivity analysis). Posterior density plots show the effect of each experimental contrast on the log-odds of a cell carrying at least one deletion or duplication event of ≥5 Mb, prior to application of the arm-weakness filter and chromosome-arm size criterion. Layout and encoding are identical to panel B. This analysis confirms that the elevated del/dup burden in SRSF2-edited (P(FC>1) = 1.000) and DNMT3A-edited cells (P(FC>1) = 0.997) is robust to the choice of size threshold, while the modest AZD7648 reduction trend observed at this threshold (P(FC<1) = 0.616) becomes less apparent after applying more stringent filters, consistent with the 5 Mb threshold retaining a higher proportion of small recurrent technical events. **(B)** Per-cell sister chromatid exchange (SCE) counts across all conditions. Each point represents one single cell; the distribution of SCE counts per condition is shown across all 961 cells from three biological replicates. SCE events were extracted from the MosaiCatcher stringent quality filters (minimum allele frequency = 5%, minimum inter-SCE distance = 500 kb). Conditions are shown on the x-axis (Ctrl, SRSF2, SRSF2 + AZD, DNMT3A, DNMT3A + AZD, Dual + AZD). **(C)** Chromatin region distribution of SV events. Stacked bar plot showing the proportion of validated SV events (del/dup and inversions combined) occurring in euchromatin or heterochromatin regions, per condition. Colours: euchromatin = light grey, heterochromatin = dark grey. **(D)** Deletion and duplication size distribution per condition after filtering steps. Violin plots show the full distribution; overlaid boxplots show median (red line) and interquartile range. Y-axis in log scale kilobases (kb). No significant differences in event size were detected between conditions (Mann-Whitney U test). **(E)** Inversion distribution per condition after filtering steps. Violin plots show the full distribution; overlaid boxplots show median (red line) and interquartile range. Y-axis in log scale kilobases (kb). No significant differences in event size were detected between conditions (Mann-Whitney U test). **(F)** Correlation between HDR integration efficiency (%) and the percentage of cells carrying at least one validated reciprocal translocation event across all edited conditions and biological replicates. Each data point represents one condition × replicate × locus combination; colour indicates the edited locus (red = SRSF2, blue = DNMT3A); filled markers indicate conditions with AZD7648, open markers indicate conditions without AZD7648; triangles indicate Dual+AZD conditions; circles indicate Control. The dashed line shows the ordinary least-squares regression fit across all non-control data points. R² and p-value are indicated. **(G)** Correlation between HDR integration efficiency (%) and the percentage of cells carrying at least one validated deletion or duplication event (stringent filter: ≥10% chromosome arm length with arm-weakness filter applied). Visual encoding as in (F). **(H)** Correlation between HDR integration efficiency (%) and the percentage of cells carrying at least one validated inversion event (arm-weakness filter with shared-control blacklist at reciprocal overlap ≥0.6). Visual encoding as in **(F)**.

**Supplementary Figure 8. Genome-wide somatic structural variant hotspot map.** Same representation as Figure 5A for all autosomes except chr2 and chr17 (chr1, chr3–chr16, chr18–chr22). Chromosomes are arranged in groups of four per row across five rows, with all six experimental conditions displayed side by side within each chromosome group to facilitate direct comparison across conditions. For each condition, deletions and duplications combined (red) and inversions (teal) are shown as individual full-span bars to the right of the shared ideogram, with colour opacity reflecting the number of locally overlapping events. Only large structural variants representing at least 10% of the corresponding chromosome arm length were retained, and events overlapping centromeric and pericentromeric regions were excluded prior to visualization. Colour opacity reflects the number of locally overlapping events, with each additional overlapping event contributing 0.20 opacity (1 event = lightest, ≥5 events = near opaque).

## Methods

### Human cord blood processing and cryopreservation

Anonymized consented human umbilical cord blood was obtained from Hôpital St-Justine and Hema Quebec, Montréal, QC, Canada. Ethics approval for the use of these cells was granted from the Comité d’éthique de la recherche clinique (CERC) of the Université de Montréal. CD34+ cells were isolated using EasySep™ Human CD34 Positive Selection Kit II (STEMCELL Technologies, Vancouver, BC, Canada) as per manufacturer instructions, and then used either directly or cryopreserved in fetal bovine serum (Gibco) and 10% dimethyl sulfoxide (BioShop). Cells were frozen slowly at -80 °C in a CoolCell (Corning) and transferred the following day to liquid nitrogen until use. On thaw, cells were rapidly warmed to 37 °C in a water bath, diluted 10x in RPMI + 10% FBS (Gibco) and spun to remove media and residual DMSO. Viable cells were counted using a hemocytometer with Trypan Blue (Gibco) and placed into culture.

### CD34+ cell culture

CD34+ cells were cultivated in StemSpan™ SFEM II media supplemented with growth factors (SCF and FLT3L at 100 ng/mL, IL3 and IL6 at 20ng/mL, Cedarlane (GenScript USA Inc)) and 35 nM UM171. Cell density was maintained below 250000 cells/mL of total media to prevent autoinhibition of the primitive cells. Cells were maintained in a humidified incubator at 37 °C with periodic viable cell counts (by hemocytometer) to ensure that the density remained within the acceptable range.

### RNP assembly

The tracrRNA and crRNA (IDT) were mixed at equimolar ratio to a final concentration of 100 μM, annealed for 5min at 95 °C and cooled to 25 °C at 0.1 °C/s. Annealed gRNA was then added to Cas9 enzyme (IDT) and incubated at room temperature for 15 min with a ratio of Cas enzyme to gRNA of 1:2.5.

### CD34+ cell editing

After 48 hours of pre-stimulation, viable cells were counted by hemocytometer. Prior to nucleofection, cells were washed once with PBS, spun down for 5 minutes at 300 g, and re-suspended in buffer P3 (Lonza) such that each well of the Nucleocuvette strip would contain 20 000 – 100 000 cells. Assembled RNP, p53 siRNA (20 fmol, Thermo id s605), and any ssODN donors (IDT) (as specified in for each experiment). Overall RNP and other additives were kept at or below 10% of the total 20 μL volume per well. Handling time between wash and nucleofection was kept within a 10-minute window. Cells were nucleofected using the Lonza 4D nucleofector device with nucleocuvette strips, using buffer setting Primary P3 and program DZ100. Following nucleofection, cells were allowed to rest for 5 minutes then added to pre-warmed wells of a 24 well plate containing media (as specified in CD34+ cell culture) supplemented with small molecules as indicated for specific experiments (AZD7648 (5µM) (Cayman Chemicals) and PolQi2 (3µM) (Cederlane, Medchem)). Cells were incubated for an additional 48 hours prior to subsequent use for nanopore sequencing. For Strand-seq experiments, cells were incubated for 6 more days for expansion in culture media, before cryopreservation and shipping for Strand-seq which was performed according to the protocols in Hanlon et al (32).

### Genomic DNA lysis and amplification for editing efficiency assessment using Sanger sequencing

At time of harvest, 5% of the cells were centrifuged at 300g for 5 min and washed once with PBS and pelleted again followed by re-suspension in a gDNA lysis buffer (50 mM Tris, 1 mM EDTA, 0.5% Tween-20 and 16 U/mL Proteinase K). Samples were incubated 1 hour at 37°C and 10 minutes at 95°C. Lysates were amplified by PCR using the Platinum Taq SuperFi II Master Mix with 0.5 μM of each primer, and lysate comprising no more than 5% of the total volume of the reaction. For SRSF2 short ssODN with silent mutations, amplification was performed with pri0077-F + pri0077-R (Supplementary methods). For DNMT3A short ssODN with silent mutations, amplification was performed with pri0245-F + pri0245-R (Supplementary methods). For all PCRs, the program was 98°C 30 s; 35× (98°C 10 s, 60°C, 10 s, 72°C 30 s); 72°C 5 min. PCR products were purified by GeneJet PCR purification, quantified by nanodrop, and 5-15 ng sent for Sanger sequencing at the IRIC Genomics core with the primer pri0003-A1 for SRSF2 donor and pri0245-F for DNMT3A donor (Supplementary methods). Integration was then quantified using ICE Analysis (Synthego Performance Analysis, 2019. v3.0. Synthego) with comparison between each edited sample back to a matched unedited control.

### Genomic amplification for nanopore sequencing

Genomic DNA from lysis step have been used for large deletions assessment at on-target sites for SRSF2 and DNMT3A. For SRSF2 large amplification, the same set of primers were used than for editing efficiency assessment for an amplicon size of 1157bp (Supplementary methods). For amplicon size of 4188bp, primer set: pri0503-F and pri0503-R were used (Supplementary methods). For DNMT3A on-target site, pri0374-F + pri0375-R have been used for amplicon size of 5137bp (Supplementary methods). For all PCRs, the program was 98°C 30 s; 30× (98°C 10 s, 60°C, 10 s, 72°C 2min30s); 72°C 5 min. PCR products were purified by GeneJet PCR purification and quantified by nanodrop.

### Editing efficiency measurement and comparison

Editing efficiency was assessed in parallel by Sanger sequencing (percentage of indel-containing reads, quantified by ICE synthego) and by the Nanopore pipeline (percentage of reads with any indel). Agreement between the two methods was evaluated by ordinary least-squares (OLS) linear regression.

### Amplicon design and read depth filtering

Two PCR amplicon sizes were used for SRSF2 across experimental batches: a 1157 bp amplicon (Batch 1) and a 4188 bp amplicon (Batch 2). DNMT3A was amplified with a 5137 bp amplicon in both batches. Because amplicon length constrains the maximum detectable deletion size, the 1157 bp amplicon systematically underestimates large deletion frequency relative to longer amplicons (Figure S6B–D). Accordingly, only the 4188 bp SRSF2 amplicon was used for inter-locus comparisons (Figure 1A–E).

### Analysis of on-target editing by nanopore sequencing

Amplicons for each of SRSF2 and DNMT3A sites were first pooled per corresponding condition at equimolar ratios based on their known length and concentration measured by Nanodrop. This allows higher multiplexing without requiring additional barcoding reagents as each amplicon will map uniquely. Each pool was then end repaired using the NEBNext® Ultra™ II End Repair/dA-Tailing Module with the addition of DNA Control Sample (Oxford Nanopore) as per manufacturer instructions, and end-repaired/A-tailed products purified using AmpureXP beads at a 1x bead ratio. Unique native barcodes were then added to each repaired/tailed amplicon pool using the Native Barcoding Kit 24 V14 (Oxford Nanopore) and NEB Blunt/TA Ligase Master Mix as per manufacturer instructions, all barcoded amplicons pooled into a single tube, and the library purified using AmpureXP beads, this time at a 0.7x bead ratio. Finally, sequencing adapters were added to the pooled library using the NEBNext® Quick Ligation Module (NEB) with the native adapter from the Native Barcoding Kit 24 V14, and this again purified with AmpureXP beads using Short Fragment Buffer (Oxford Nanopore) instead of 80% ethanol, all as per manufacturer instructions. Final concentration was determined by Nanodrop and 20 fmol loaded onto a Flongle Flow Cell (R10.4.1) with a minION sequencing device (MIN-101B) with Flongle adapter (all from Oxford Nanopore). Samples were run with MinKNOW, and base-calling executed on Super-Accurate mode.

Following base-calling, samples were analyzed using a custom two-stage pipeline (https://github.com/djhfknapp/Nanopore-Large-Deletion-Analysis). For each sample, FASTQ files were pooled and aligned to the expected amplicon reference sequences derived from GRCh38 using minimap2 with splice-aware alignment parameters optimized for large deletion detection (-ax splice --splice-flank=no -O 6,24 --secondary=no). Alignments were converted to BAM format, filtered to retain only primary alignments with mapping quality ≥10 (samtools view -q 10 -F 2048), sorted, and indexed using samtools. Target sites were supplied as a tab-delimited file containing the expected guide sequence for each amplicon. The target sequence was localized within each reference amplicon by searching both forward and reverse-complement orientations, and genomic coordinates corresponding to the target site were recorded for downstream analyses.

Large deletion analysis was performed prior to variant calling. Each primary alignment was interrogated at the CIGAR-string level and all deletion (D) and skipped-region (N) operations were examined. A deletion was classified as a large deletion if: (i) its size was at least 50 bp, (ii) at least 20 aligned bases were present on both sides of the deletion, (iii) the deletion overlapped the predicted target site, and (iv) the read contained no more than one qualifying large deletion event. Reads failing these criteria were excluded from large deletion analyses. For each qualifying event, deletion coordinates, deletion size, affected reference sequence, and read identifier were recorded. Summary statistics including total reads, reads containing large deletions, deletion frequencies, and deletion size distributions were generated for each amplicon.

To prevent large structural events from confounding small variant analysis, reads containing large deletions were removed prior to variant calling. A filtered BAM file containing only reads without qualifying large deletions was generated and indexed. Small variants were then called on this filtered BAM using bcftools mpileup with parameters -Ov -Q 16 -a FORMAT/DP,FORMAT/AD -d 10000 --min-MQ 10. Variant calls were summarized on a per-position basis by extracting reference depth, alternate allele depth, and allele frequencies from the resulting VCF file. For editing outcome quantification, predicted target sites were identified within each amplicon and variants were classified as substitutions, insertions, or deletions if they originated within or extended into the target region. Variants supported by fewer than three reads or representing less than 0.1% of reads at a given site were excluded. Per-site editing summaries were generated by calculating the percentage of reads containing substitutions, insertions, or deletions overlapping the target region. Reference allele frequencies were calculated as the proportion of reads not containing a qualifying variant within the target site. For visualization, coverage profiles were generated for each reference sequence and per-read deletion maps were produced showing the position and extent of large deletions relative to the target site.

### Bayesian mixed-effect model for percentage of large deletions and average size deletions

A quality control filter was applied prior to modelling: any Gene×Replicate combination in which the unedited control sample had fewer than 1,000 total reads was excluded entirely for that gene in that replicate, removing 12 observations and leaving 60 samples. For the deletion size analysis, SRSF2 samples sequenced on the shorter 1,157 bp amplicon (Batch 1, n = 21 rows) were additionally excluded, as this amplicon is not comparable to the longer 4,188 bp amplicon used in Batch 2 for deletion size estimation. The deletion size analysis therefore used 42 observations (36 with Sex recorded).

Two binary predictors were derived from the Condition and Gene columns: SRSF2 (1 = SRSF2-edited condition, 0 = unedited SRSF2 control or any DNMT3A row) and DNMT3A (1 = DNMT3A-edited condition, 0 = unedited DNMT3A control or any SRSF2 row). These two indicators are never simultaneously equal to 1. Drug treatment was encoded as AZD7648 (1 = AZD or AZD7648+PolQi2 condition) and PolQi2 (1 = PolQi or AZD7648+PolQi2 condition). Biological sex was encoded as a factor with Female as the reference level. Both models were fitted including Sex as a covariate (primary, n = 48 and n = 36 for frequency and size respectively).

#### Model 1: Large deletion frequency (Beta regression)

The proportion of Nanopore reads containing large deletions (reads_with_large_del / total_reads) was modelled using a Bayesian mixed-effects Beta regression, implemented in the brms package (version 2.23.0) in R (version 4.4.3) with Stan as the MCMC backend. Beta regression was chosen because the outcome is a proportion strictly bounded between 0 and 1, and models data on the logit scale without requiring a sequencing depth offset, since the proportion already accounts for variation in total read depth across samples. All values satisfied the strict (0, 1) constraint required by the Beta likelihood (minimum observed proportion = 0.0004, maximum = 0.60).

The model formula was:

prop ∼ (SRSF2 + DNMT3A) × AZD7648 × PolQi2 + Sex + (1 | Replicate)

The unedited control (Ctrl) served as the reference for the editing terms (SRSF2 and DNMT3A), while the editing-only condition (CRISPR without AZD7648 and PolQi2) served as the reference for the drug effect terms, such that AZD7648 and PolQi2 coefficients reflect changes in large deletion rates on top of CRISPR editing alone. Replicate was included as a random intercept to account for repeated measurements across cord blood donors. Weakly informative priors were specified for all parameters. Fixed effects received normal(0, 2) priors on the logit scale. The intercept received a Student-t(3, −4, 3) prior, centered near the logit-transformed baseline deletion rate in unedited controls (∼0.1–0.2%, logit ≈ −6.9). Random effect standard deviations received half-normal(0, 1) priors. The Beta precision parameter φ received a gamma(0.01, 0.01) prior.

#### Model 2: Average large deletion size (Gaussian linear model)

The average size of large deletions per sample (avg_del_size, in base pairs) was modelled using a Bayesian mixed-effects Gaussian linear model with an identity link function, using the same brms framework as previously. A Gaussian likelihood was chosen because the distribution of average deletion sizes across samples was approximately normal (skewness = 0.18, range 256–3,719 bp), and no transformation was required. Coefficients are directly interpretable in base pairs.

The model formula was:

avg_del_size ∼ (SRSF2 + DNMT3A) × AZD7648 × PolQi2 + Sex + (1 | Replicate)

The reference structure and random effects specification were identical to Model 1. Weakly informative priors were specified on the base pair scale: fixed effects received normal(0, 500) priors, allowing shifts of up to ∼1,000 bp (2 SD); the intercept received a Student-t(3, 1500, 500) prior centred near the observed overall mean of 1,482 bp; the residual standard deviation (sigma) received a half-Student-t(3, 0, 500) prior; and random effect standard deviations received half-normal(0, 300) priors.

#### Model fitting, convergence, and posterior inference

Both models were fitted using four parallel Markov chain Monte Carlo (MCMC) chains, each with 3,000 iterations including 1,000 warmup draws, yielding 8,000 post-warmup posterior draws in total. The No-U-Turn Sampler (NUTS) was used with adapt_delta = 0.99 and max_treedepth = 15 to ensure stable sampling. Convergence was assessed using the potential scale reduction factor (Ř-hat), with all parameters achieving Ř-hat ≤ 1.01, and bulk and tail effective sample sizes exceeding 2,000 for all parameters.

Drug effects were evaluated as composite posterior contrasts combining the shared drug (AZD7648 and polQi2) main effect and the gene-specific interaction term. For example, the total effect of AZD7648 in SRSF2-edited cells was computed as b_AZD7648 + b_SRSF2:AZD7648, and equivalently for DNMT3A. The total effect of the AZD7648 + PolQi2 combination versus editing alone was computed by summing all relevant drug coefficients and their interactions. All contrasts were evaluated using the hypothesis() function in brms, which computes posterior probabilities (PP) and evidence ratios (Bayes factors) directly from the posterior draws. For Model 1, results are reported on the odds ratio scale (exp(logit coefficient)), where values above 1 indicate an increase and values below 1 indicate a decrease in the proportion of reads with large deletions. For Model 2, results are reported directly in base pairs. All analyses were conducted using R version 4.4.3, brms version 2.23.0, and Stan via the rstan interface.

### MMEJ scoring and assessment

To assess whether microhomology-mediated end joining (MMEJ) contributes to CRISPR-induced deletions at the DNMT3A R882 and SRSF2 P95 loci using a canonical microhomology threshold, we scanned all Oxford Nanopore sequencing reads from PCR amplicons spanning the DNMT3A cut site for deletion events bearing microhomology ≥ 5 bp at the breakpoint junction. Unlike the primary analysis which focused on large deletions (≥ 50 bp) identified from pre-filtered BAM files, this analysis was performed on all reads in the main alignment file, including those carrying small deletions (≥ 10 bp), to maximise sensitivity. For each read, deletion events were extracted from the CIGAR string (D and N operations), and microhomology was scored by comparing the reference sequence flanking each breakpoint. Read-level junction confirmation was performed using a CIGAR-aware approach: the exact query positions flanking the deletion were identified using get_aligned_pairs(), and the junction sequence was inspected to verify retention of a single copy of the microhomology, consistent with MMEJ repair. Only deletions overlapping the CRISPR cut site and passing read-level confirmation were classified as MMEJ events. MMEJ frequency was expressed as the percentage of total reads at the locus bearing a confirmed ≥ 5 bp microhomology at the cut site junction. Analysis was performed across two independent batches comprising 7 biological replicates for the no molecule reference and AZD7648 conditions, and 3 replicates for the PolQi2 and PolQi2+AZD7648 conditions.

### Strand Seq library preparation and sequencing

Single-cell Strand-seq libraries were prepared using a streamlined version of the established Strand-seq protocol(32,33). HSPCs were cultured for 24 h in the presence of bromodeoxyuridine (BrdU) and nuclei with BrdU incorporation in the G1 phase of the cell cycle were sorted by fluorescence-activated cell sorting (FACS) as described previously (32). Single nuclei were dispensed into individual wells of an open 72 × 72 well nanowell array and treated with heat-labile protease, followed by digestion of DNA with the restriction enzymes AluI and HpyCH4V (NEB). Fragments were A-tailed, ligated to forked Illumina adapters, UV-treated to ablate the BrdU-substituted strand, and PCR-amplified with indexed primers. Libraries were pooled and cleaned with AMPure XP beads, and library fragments between 300 and 700 bp were gel-purified before PE75 sequencing on NextSeq 550 or the AVITI(33). Reads were aligned to the human reference genome (hg38) and processed through the MosaiCatcher pipeline or quality control, normalization, and structural variant calling(34). Libraries passing quality thresholds were retained for downstream analysis. For a comprehensive overview of Strand-seq principles and applications, see(21)

### Strand-Seq data processing for translocation assessment

Pseudo-bulk BAM preparation: to increase statistical power for structural rearrangement detection, single-cell BAM files were merged per condition using SAMtools merge (v1.22.1) with 8 parallel threads. All cells passing MosaiCatcher quality control were included, yielding pseudo-bulk BAMs representing 189 cells (control), 175 cells (SRSF2 no-AZD7648), 82 cells (SRSF2 + AZD7648), 174 cells (DNMT3A no-AZD7648), 84 cells (DNMT3A + AZD7648), and 297 cells (dual + AZD7648). Merged BAMs were indexed using SAMtools index.

Genome-wide reciprocal rearrangement detection: inter-chromosomal structural rearrangements were detected from single-cell BAM files using a custom Python pipeline leveraging pysam (v0.23.3) (https://github.com/Fanny-Mei/scReciprocalTransloc). For each cell, a two-pass strategy. In the first pass, all discordant read pairs (reads mapping to different canonical autosomes) were enumerated across all chromosome pairs, applying minimum mapping quality (MAPQ ≥30) and excluding duplicate, secondary, and supplementary alignments. Chromosome pairs with at least two discordant reads on each side (chrA→chrB and chrB→chrA) were retained as candidates for the second pass. In the second pass, candidate chromosome pairs underwent validation by three additional filters applied simultaneously: (i) strand orientation consistency (≥65% majority strand on each side), (ii) positional clustering (70% of reads within a 5 Mb window), and (iii) confirmed reciprocal support on both sides. A chromosome pair in a given cell was classified as a validated rearrangement only if all four criteria were simultaneously satisfied. Breakpoint positions were recorded as the median, minimum, and maximum genomic coordinates of supporting reads on each side.

CRISPR cut site proximity annotation: validated rearrangements were annotated for proximity to CRISPR guide RNA cut sites. The SRSF2 P95H guide RNA (spacer: CGGCTGTGGTGTGAGTCCGG) targets chr17:78,015,826 and the DNMT3A R882H guide RNA (spacer: GCAGTCTCTGCCTCGCCAAG) targets chr2:25,654,947 in the hg38 reference genome, as determined by exact sequence matching using SAMtools faidx. Rearrangements were flagged as near-cut if the median breakpoint position on either chromosome fell within 5 Mb of the respective cut site.

### Bayesian mixed-effects statistical model for translocations

To compare rearrangement rates across conditions while accounting for replicate variability and the factorial experimental design, we fitted a Bayesian mixed-effects Poisson model using brms (v2.23.0) in R (v4.4.3). Rather than treating dual editing as an independent condition, the model encodes each condition as a combination of three binary predictors, SRSF2 editing, DNMT3A editing, and AZD7648 treatment, allowing dual editing to emerge naturally as the interaction between the two editing terms. The model formula was:

*n_events ∼ srsf2 * dnmt3a * azd7c48 + offset(log(n_cells)) + (1 | replicate)*

which expands to all main effects and pairwise interactions between the three binary predictors. For each observation I (one condition-replicate combination), the total number of validated rearrangement events Yi was modelled as:

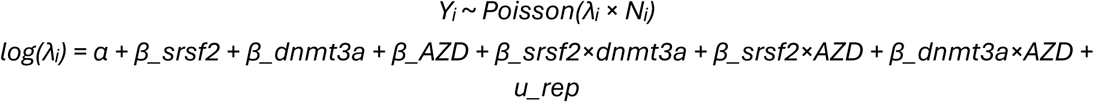

where λᵢ is the per-cell event rate, Ni is the number of cells (entered as an offset on the log scale to model rates rather than counts), α is the global intercept, β_srsf2 and β_dnmt3a capture the main effects of editing at each locus, β_AZD captures the main effect of AZD7648 treatment, and the three pairwise interaction terms capture condition specific modulation of these effects. Critically, the β_srsf2xdnmt3a term directly tests whether dual editing causes more rearrangements than the additive expectation from the two individual editing effects: a positive and credible coefficient indicates synergy, whereas a coefficient spanning zero indicates additivity. The replicate random intercept u_rep ∼Normal (0, σ_rep) accounts for batch-to-batch variability across three biological replicates.

Weakly informative priors were used throughout: intercept Normal(-2.3, 1.0) (centered on log (0.1) ≈ -2.3, corresponding to a prior expectation of ∼0.1 events per cell), fixed effects Normal(0, 0.5), and replicate standard deviation half-Normal(0, 0.5). Marginal condition-level posterior rates were derived from posterior draws of the linear predictor evaluated at u_rep = 0, corresponding to the population-average rate. Posterior distributions were estimated using the No-U-Turn Sampler (NUTS) with 4 chains of 4000 draws each (2000 warmup steps, target acceptance rate adapt_delta = 0.95). Convergence was confirmed by the Gelman-Rubin statistic (R ≤ 1.01 for all parameters). Posterior probabilities P(λ_A > λ_B) were computed as the proportion of posterior samples where condition A exceeded condition B.

### Single-cell Strand-seq SV calling

Single-cell Strand-seq libraries were prepared from 3 replicates across six experimental conditions: Ctrl, *SRSF2* no mol, *SRSF2* + AZD, *DNMT3A* no mol, *DNMT3A* + AZD, and Dual + AZD (simultaneous editing of both loci with AZD7648). A total of 961 single-cell libraries passed quality control performed using Ashley’s pipeline. Structural variants were called using the MosaiCatcher pipeline v2 (Weber et al., 2023; hg38 assembly, default parameters, window size = 200 kb)(34). Analysis was restricted to autosomes (chr1–22). SV calls were classified into four categories based on the sv_call_name field: deletion, duplication (including inverted duplications), inversion, and complex events. Donor sex was determined computationally from the chrY segmentation strand state (WC = female; WW or CC = male).

### Event filtering strategy for deletion/duplication and inversion analyses

SV calls produced by the MosaiCatcher pipeline were post-processed independently for deletion/duplication and inversion event classes, each using a dedicated filtering strategy designed to minimise technical noise while preserving genuine somatic rearrangements.

#### Common pre-filters applied to all event classes

Prior to class-specific filtering, the following exclusions were applied universally: (i) events assigned to the WC strand class were excluded, as the WC configuration precludes reliable haplotype-resolved SV calling; (ii) for duplication calls, CW dup_hom events were excluded; (iii) events with more than 50% overlap with annotated centromeric regions were excluded to avoid centromere-driven artefacts.

#### Deletion and duplication — pre-filter analysis (5 Mb threshold)

For the sensitivity analysis, adjacent same-cell, same-arm, same-type calls were first merged if separated by less than 200 kb. Merged events were retained only if their final size was ≥5 Mb, corresponding to the window resolution used in the MosaiCatcher segmentation. No chromosome-arm percentage threshold, arm-weakness filter, shared-control blacklist, control-like cell exclusion, or chromosome-spread cell exclusion was applied at this stage. This yielded a total of 1,295 retained events across 191 event-positive cells: Control 129 events in 26/186 cells (14.0%), SRSF2 685 events in 48/166 cells (28.9%), SRSF2+AZD 29 events in 14/78 cells (18.0%), DNMT3A 91 events in 43/170 cells (25.3%), DNMT3A+AZD 64 events in 12/77 cells (15.6%), and Dual+AZD 297 events in 48/284 cells (16.9%).

#### Deletion and duplication — final analysis (≥10% arm, arm-weakness filter)

Given the high residual event burden at the 5 Mb threshold, a more stringent criterion was applied for the primary analysis. Events were retained only if they spanned at least 10% of the length of the chromosome arm on which they were called. An arm-weakness filter was then applied: a chromosome arm was excluded from all non-control cells if more than one control cell carried a candidate event on that arm and at least one non-control cell also carried a candidate event on that arm. Following arm-weakness filtering, two downstream cell-level exclusions were applied: (i) non-control cells in which more than one retained event overlapped a pre-blacklist control event at reciprocal overlap ≥0.5 were removed as control-like cells; (ii) cells with retained events on more than five distinct chromosomes were excluded as chromosome-spread cells. This yielded a total of 154 retained events across 72 event-positive cells: Control 13 events in 6/186 cells (3.2%), SRSF2 55 events in 16/166 cells (9.6%), SRSF2+AZD 7 events in 5/78 cells (6.4%), DNMT3A 19 events in 13/170 cells (7.6%), DNMT3A+AZD 8 events in 5/77 cells (6.5%), and Dual+AZD 52 events in 27/284 cells (9.5%).

#### Inversion events

Inversion calls were filtered independently of del/dup events. No chromosome-arm size threshold was applied. Events with any centromeric overlap (>0%) were excluded. WC strand-class calls were excluded as above. A shared-control blacklist was applied: inversion events were removed if they overlapped a control-cell inversion call at reciprocal overlap ≥0.6, reflecting the higher recurrence rate of inversion artefacts at specific loci. An arm-weakness filter was then applied using the same logic as for del/dup: a chromosome arm was excluded if more than one control cell carried an inversion candidate on that arm and at least one non-control cell also had a candidate on that arm. This led to the exclusion of seven arms: chr1p, chr2p, chr2q, chr5q, chr6q, chr9p, and chr20q. No control-like cell or chromosome-spread cell exclusion was applied to inversions. Final retained counts were: Control 13/186 cells (7.0%), SRSF2 23/166 (13.9%), SRSF2+AZD 13/78 (16.7%), DNMT3A 25/170 (14.7%), DNMT3A+AZD 16/77 (20.8%), Dual+AZD 55/284 (19.4%).

### Chromatin region enrichment analysis

Genomic region classes were defined from the UCSC hg38 cytoBandIdeo annotation and collapsed into two broad categories: euchromatin (gneg, gpos25) and heterochromatin (gpos50, gpos75, gpos100, gvar). SV events were assigned to regions based on maximum base-pair overlap. The proportion of validated SV events (del/dup and inversions combined) occurring in each chromatin class was computed per condition and visualized as stacked bar plots. No statistical testing was applied to the chromatin distribution, as the analysis was intended to describe the overall accessibility bias of retained events rather than to test condition-specific enrichment.

### Bayesian mixed-effects cell-rate models for del/dup and inversion events

To assess the effects of CRISPR editing, AZD7648 treatment, and donor sex on the probability of a cell carrying at least one structural variant event, we fitted three independent Bayesian mixed-effects binary outcome models: one for del/dup events prior to stringent filtering (5 Mb size threshold), one for del/dup events after final filtering (≥10% chromosome arm, arm-weakness filter applied), and one for inversion events (arm-weakness filter with shared-control blacklist at reciprocal overlap ≥0.6). All models were implemented in brms (v2.23.0) on R (v4.4.3) with Stan as the MCMC backend. The response variable was a binary indicator of whether each cell carried at least one retained event of the relevant SV class (0 = event-negative, 1 = event-positive). The linear predictor on the log-odds scale was:

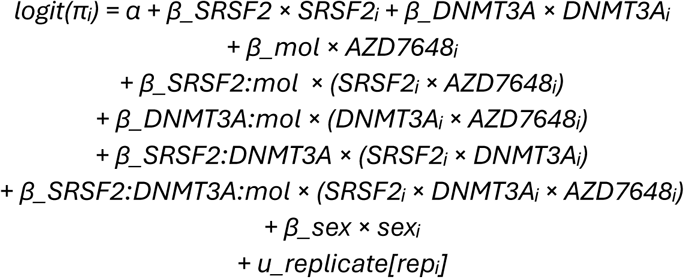

Population-level (fixed) effects included: SRSF2 editing status (0/1), DNMT3A editing status (0/1), and AZD7648 treatment (0/1), each encoded as binary indicators with unedited/untreated cells as the reference level. Dual-edited cells were encoded as SRSF2=1 and DNMT3A=1 simultaneously, such that their total effect decomposes into individual editing contributions (β_SRSF2 + β_DNMT3A) plus a synergy term (β_SRSF2:DNMT3A) capturing any excess above additive expectation on the log-odds scale. The three-way interaction term β_SRSF2:DNMT3A:mol tests whether the dual-editing synergy is further modified by AZD7648 treatment. Donor sex was included as a binary population-level covariate (female = reference). Group-level (random) intercepts were included per biological replicate to account for batch effects across cord blood donors.

Weakly informative priors were specified as: normal(0, 1.5) for population-level coefficients, Student-t(3, 0, 2.5) for the intercept, and exponential(1) for the random-effect standard deviation. Each model was run with 4 chains of 4,000 iterations (1,000 warm-up), with adapt_delta = 0.95 and max_treedepth = 12. Model effects are reported as posterior probabilities P(β > 0) or P(β < 0) as appropriate for the direction of the contrast, with effect sizes expressed as fold-change (FC = exp(β)) of the posterior median.

### Sister chromatid exchange analysis

SCE events were extracted from the MosaiCatcher stringent quality filter (minimum allele frequency = 5%) using the total_sce column from the per-cell stats-merged.tsv output. SCEs are detected in Strand-seq as cell-private strand-state switches, mid-chromosome changes between Watson/Crick ground states without read-depth changes, modelled separately from SVs by Mosaicatcher using a minimum inter-SCE distance of 500 kb. Per-cell SCE counts were compiled across all 961 cells.

### Hotspot analysis and ideogram visualization

Structural variant events were filtered prior to visualisation by two sequential criteria: events overlapping centromeric or pericentromeric regions (as defined in the Chromatin region enrichment analysis section) were excluded, and only events representing at least 10% of the corresponding chromosome arm length were retained, as determined from the UCSC hg38 cytoBandIdeo annotation. Chromosome arm lengths were computed separately for the p and q arms using the centromere (acen) band boundaries as the delimiter, and each event was assigned to an arm based on its midpoint position. Chromosomal ideograms were rendered in Python using standard Giemsa cytoband colours (gneg = white through gpos100 = near black; centromere = deep purple). For each chromosome group, a single shared ideogram was displayed on the left, followed by two sets of six condition-specific SV tracks: deletions and duplications combined (red) and inversions (teal). Inverted duplications were merged into the duplication category prior to visualisation. Complex rearrangements were excluded from the ideogram visualisation due to insufficient event counts after filtering. Each SV event is represented as a bar spanning its full genomic coordinates. Colour opacity reflects the local density of overlapping events, with each additional overlapping event contributing 0.20 opacity (minimum 0.20 for a single event, capped at 0.95 for five or more overlapping events). The genomic positions of the DNMT3A (chr2:25,285,000) and SRSF2 (chr17:74,733,000) loci were annotated on the respective chromosome ideograms as horizontal tick lines (DNMT3A = blue, SRSF2 = red). All six experimental conditions are displayed side by side for each chromosome to facilitate direct visual comparison. All analyses were performed in Python 3.12 using pandas, matplotlib, scipy, and statsmodels.

## References

1. Lee BC, Lozano RJ, Dunbar CE. Understanding and overcoming adverse consequences of genome editing on hematopoietic stem and progenitor cells. Mol Ther. 2021 Nov;29(11):3205–18. doi:10.1016/j.ymthe.2021.09.001

2. Robert F, Barbeau M, Éthier S, Dostie J, Pelletier J. Pharmacological inhibition of DNA-PK stimulates Cas9-mediated genome editing. Genome Med. 2015 Dec;7(1):93. doi:10.1186/s13073-015-0215-6

3. Hunt JMT, du Rand A, Verdon D, Clemance L, Loef E, Malhi C, et al. Enhanced HDR-mediated correction of heterozygous COL7A1 mutations for recessive dystrophic epidermolysis bullosa. Mol Ther Nucleic Acids. 2025 Mar 11;36(1):102472. doi:10.1016/j.omtn.2025.102472 PubMed PMID: 40027886; PubMed Central PMCID: PMC11872078.

4. Selvaraj S, Feist WN, Viel S, Vaidyanathan S, Dudek AM, Gastou M, et al. High-efficiency transgene integration by homology-directed repair in human primary cells using DNA-PKcs inhibition. Nat Biotechnol. 2024 May;42(5):731–44. doi:10.1038/s41587-023-01888-4

5. Cloarec-Ung FM, Beaulieu J, Suthananthan A, Lehnertz B, Sauvageau G, Sheppard HM, et al. Near-perfect precise on-target editing of human hematopoietic stem and progenitor cells. eLife. 2024 Jun 3;12:RP91288. doi:10.7554/eLife.91288.3

6. Pugliano CM, Berger M, Ray RM, Sapkos K, Wu B, Laird A, et al. DNA-PK inhibition enhances gene editing efficiency in HSPCs for CRISPR-based treatment of X-linked hyper IgM syndrome. Mol Ther - Methods Clin Dev. 2024 Sep;32(3):101297. doi:10.1016/j.omtm.2024.101297

7. Wimberger S, Akrap N, Firth M, Brengdahl J, Engberg S, Schwinn MK, et al. Simultaneous inhibition of DNA-PK and Polϴ improves integration efficiency and precision of genome editing. Nat Commun. 2023 Aug 14;14(1):4761. doi:10.1038/s41467-023-40344-4

8. Hunt JMT, Samson CA, Rand A du, Sheppard HM. Unintended CRISPR-Cas9 editing outcomes: a review of the detection and prevalence of structural variants generated by gene-editing in human cells. Hum Genet. 2023 Jun 1;142(6):705–20. doi:10.1007/s00439-023-02561-1

9. Little SR, Rahbari N, Hajiaghayi M, Gholizadeh F, Cloarec-Ung FM, Phillips J, et al. A Digital Microfluidic Platform for the Microscale Production of Functional Immune Cell Therapies. Anal Chem. 2025 Jun 3;97(21):11026–34. doi:10.1021/acs.analchem.4c06911

10. Haider S, Mussolino C. Fine-Tuning Homology-Directed Repair (HDR) for Precision Genome Editing: Current Strategies and Future Directions. Int J Mol Sci. 2025 Apr 25;26(9):4067. doi:10.3390/ijms26094067

11. Gavande NS, VanderVere-Carozza PS, Pawelczak KS, Mendoza-Munoz P, Vernon TL, Hanakahi LA, et al. Discovery and development of novel DNA-PK inhibitors by targeting the unique Ku–DNA interaction. Nucleic Acids Res. 2020 Nov 18;48(20):11536–50. doi:10.1093/nar/gkaa934

12. Fok JHL, Ramos-Montoya A, Vazquez-Chantada M, Wijnhoven PWG, Follia V, James N, et al. AZD7648 is a potent and selective DNA-PK inhibitor that enhances radiation, chemotherapy and olaparib activity. Nat Commun. 2019 Nov 7;10(1):5065. doi:10.1038/s41467-019-12836-9

13. Ramsden DA, Carvajal-Garcia J, Gupta GP. Mechanism, cellular functions and cancer roles of polymerase-theta-mediated DNA end joining. Nat Rev Mol Cell Biol. 2022 Feb;23(2):125–40. doi:10.1038/s41580-021-00405-2

14. Wyatt DW, Feng W, Conlin MP, Yousefzadeh MJ, Roberts SA, Mieczkowski P, et al. Essential Roles for Polymerase θ-Mediated End Joining in the Repair of Chromosome Breaks. Mol Cell. 2016 Aug 18;63(4):662–73. doi:10.1016/j.molcel.2016.06.020 PubMed PMID: 27453047; PubMed Central PMCID: PMC4992412.

15. Boutin J, Fayet S, Marin V, Bergès C, Riandière M, Toutain J, et al. Single-cell multiplex approaches deeply map ON-target CRISPR-genotoxicity and reveal its mitigation by palbociclib and long-term engraftment. Nat Commun. 2026 Jan 10;17(1):1429. doi:10.1038/s41467-025-68177-3

16. Cullot G, Aird EJ, Schlapansky MF, Yeh CD, Van De Venn L, Vykhlyantseva I, et al. Genome editing with the HDR-enhancing DNA-PKcs inhibitor AZD7648 causes large-scale genomic alterations. Nat Biotechnol. 2024 Nov 27. doi:10.1038/s41587-024-02488-6

17. Dacquay LC, Antoniou P, Mentani A, Selfjord N, Mårtensson H, Hsieh PP, et al. Dual inhibition of DNA-PK and Polϴ boosts precision of diverse prime editing systems. Nat Commun. 2025 May 8;16(1):4290. doi:10.1038/s41467-025-59708-z

18. Jeong H, Grimes K, Rauwolf KK, Bruch PM, Rausch T, Hasenfeld P, et al. Functional analysis of structural variants in single cells using Strand-seq. Nat Biotechnol. 2023 Jun;41(6):832–44. doi:10.1038/s41587-022-01551-4

19. Falconer E, Hills M, Naumann U, Poon SSS, Chavez EA, Sanders AD, et al. DNA template strand sequencing of single-cells maps genomic rearrangements at high resolution. Nat Methods. 2012 Nov;9(11):1107–12. doi:10.1038/nmeth.2206 PubMed PMID: 23042453; PubMed Central PMCID: PMC3580294.

20. Sanders AD, Meiers S, Ghareghani M, Porubsky D, Jeong H, van Vliet MACC, et al. Single-cell analysis of structural variations and complex rearrangements with tri-channel processing. Nat Biotechnol. 2020 Mar;38(3):343–54. doi:10.1038/s41587-019-0366-x PubMed PMID: 31873213; PubMed Central PMCID: PMC7612647.

21. Hanlon VCT, Lansdorp PM. Strand-seq and the future of personalized genomics. Nat Genet. 2026 May;58(5):995–1004. doi:10.1038/s41588-026-02548-4 PubMed PMID: 41882415.

22. Amarasinghe SL, Su S, Dong X, Zappia L, Ritchie ME, Gouil Q. Opportunities and challenges in long-read sequencing data analysis. Genome Biol. 2020 Dec;21(1):30. doi:10.1186/s13059-020-1935-5

23. Samuelson C, Radtke S, Zhu H, Llewellyn M, Fields E, Cook S, et al. Multiplex CRISPR/Cas9 genome editing in hematopoietic stem cells for fetal hemoglobin reinduction generates chromosomal translocations. Mol Ther - Methods Clin Dev. 2021 Dec;23:507–23. doi:10.1016/j.omtm.2021.10.008

24. Rayner E, Durin MA, Thomas R, Moralli D, O’Cathail SM, Tomlinson I, et al. CRISPR-Cas9 Causes Chromosomal Instability and Rearrangements in Cancer Cell Lines, Detectable by Cytogenetic Methods. CRISPR J. 2019 Dec 1;2(6):406–16. doi:10.1089/crispr.2019.0006

25. Porubsky D, Höps W, Ashraf H, Hsieh P, Rodriguez-Martin B, Yilmaz F, et al. Recurrent inversion polymorphisms in humans associate with genetic instability and genomic disorders. Cell. 2022 May;185(11):1986–2005.e26. doi:10.1016/j.cell.2022.04.017

26. Sanders AD, Hills M, Porubský D, Guryev V, Falconer E, Lansdorp PM. Characterizing polymorphic inversions in human genomes by single-cell sequencing. Genome Res. 2016 Nov;26(11):1575–87. doi:10.1101/gr.201160.115

27. Scrimieri F, Calhoun ES, Patel K, Gupta R, Huso DL, Hruban RH, et al. FAM190A Rearrangements Provide a Multitude of Individualized Tumor Signatures and Neo-antigens in Cancer. Oncotarget. 2011 Feb 28;2(1–2):69–75. doi:10.18632/oncotarget.220

28. Fungtammasan A, Walsh E, Chiaromonte F, Eckert KA, Makova KD. A genome-wide analysis of common fragile sites: What features determine chromosomal instability in the human genome? Genome Res. 2012 Jun;22(6):993–1005. doi:10.1101/gr.134395.111

29. Durkin SG, Glover TW. Chromosome Fragile Sites. Annu Rev Genet. 2007 Dec 1;41(1):169–92. doi:10.1146/annurev.genet.41.042007.165900

30. Huh J, Tiu RV, Gondek LP, O’Keefe CL, Jasek M, Makishima H, et al. Characterization of chromosome arm 20q abnormalities in myeloid malignancies using genome-wide single nucleotide polymorphism array analysis. Genes Chromosomes Cancer. 2010 Apr;49(4):390–9. doi:10.1002/gcc.20748

31. Machiela MJ, Zhou W, Caporaso N, Dean M, Gapstur SM, Goldin L, et al. Mosaic chromosome 20q deletions are more frequent in the aging population. Blood Adv. 2017 Feb 14;1(6):380–5. doi:10.1182/bloodadvances.2016003129

32. Hanlon VCT, Chan DD, Hamadeh Z, Wang Y, Mattsson CA, Spierings DCJ, et al. Construction of Strand-seq libraries in open nanoliter arrays. Cell Rep Methods. 2022 Jan 24;2(1):100150. doi:10.1016/j.crmeth.2021.100150

33. Porubsky D, Dashnow H, Sasani TA, Logsdon GA, Hallast P, Noyes MD, et al. Human de novo mutation rates from a four-generation pedigree reference. Nature. 2025 Jul;643(8071):427–36. doi:10.1038/s41586-025-08922-2 PubMed PMID: 40269156; PubMed Central PMCID: PMC12240836.

34. Weber T, Cosenza MR, Korbel J. MosaiCatcher v2: a single-cell structural variations detection and analysis reference framework based on Strand-seq. Alkan C, editor. Bioinformatics. 2023 Nov 1;39(11):btad633. doi:10.1093/bioinformatics/btad633

