## Supplemental Figures for "Genome-wide structural variant landscape following HDR-enhanced CRISPR editing in human hematopoietic stem and progenitor cells"

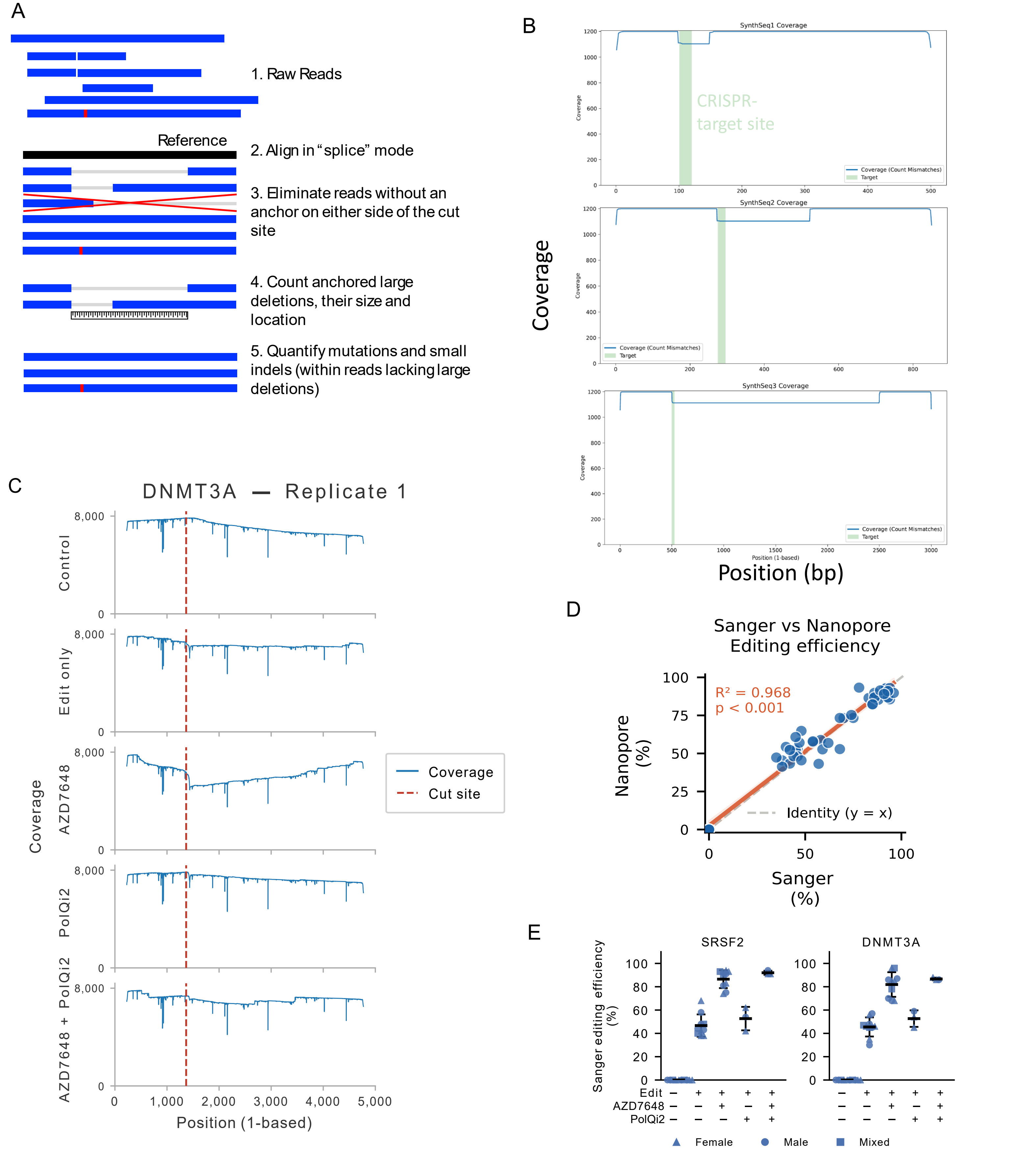


**Supplementary Figure 1. Pipeline validation for large deletion detection at on-target site. (A)** Schematic overview of the Nanopore long-read sequencing pipeline for large deletion detection. Reads are aligned to the reference amplicon and required to anchor on both sides of the targeted cut site (bilateral anchoring). Reads that do not span the full amplicon are discarded prior to quantification. Reads with an aligned deletion exceeding 50 bp relative to the reference are classified as large-deletion reads. The pipeline simultaneously reports on-target editing efficiency as the percentage of reads carrying any indel at the cut site. **(B)** Validation of pipeline sensitivity across deletion sizes using a synthetic dataset. In silico-generated deletion alleles spanning the range of detectable deletion sizes were introduced at defined frequencies and processed through the pipeline, confirming accurate detection and quantification without systematic size-dependent bias. **(C)** Coverage profile across the PCR amplicon at a representative locus, illustrating the drop in read coverage toward the 3' end characteristic of truncated Nanopore reads. These truncated reads, which cannot be assigned a true deletion size, are removed prior to large deletion quantification by the bilateral anchoring filter, ensuring that coverage drop-off artifacts do not inflate large deletion estimates. **(D)** Correlation between editing efficiency measured by Sanger sequencing (ICE analysis) and by the Nanopore pipeline across all replicates and conditions ($R^2 = 0.968$), validating the Nanopore pipeline as a reliable substitute for Sanger-based editing efficiency measurement. Each data point represents one replicate × condition × locus measurement; the solid line shows the ordinary least squares (OLS) regression fit; the shaded band represents the 95% confidence interval; the dashed line indicates the identity line (slope = 1). **(E)** Sanger sequencing-based editing efficiency (%) across all conditions at the SRSF2 and DNMT3A loci. Each data point represents one biological replicate; horizontal bars indicate the mean; error bars represent ±1 SD. Marker shape corresponds to donor sex (triangle = female, circle = male, square = mixed). AZD7648 substantially increased editing efficiency at both loci (~85–90% vs. ~47% for the p53 siRNA reference), while PolQi2 alone or in combination with AZD7648 did not substantially alter editing efficiency relative to AZD7648 alone.

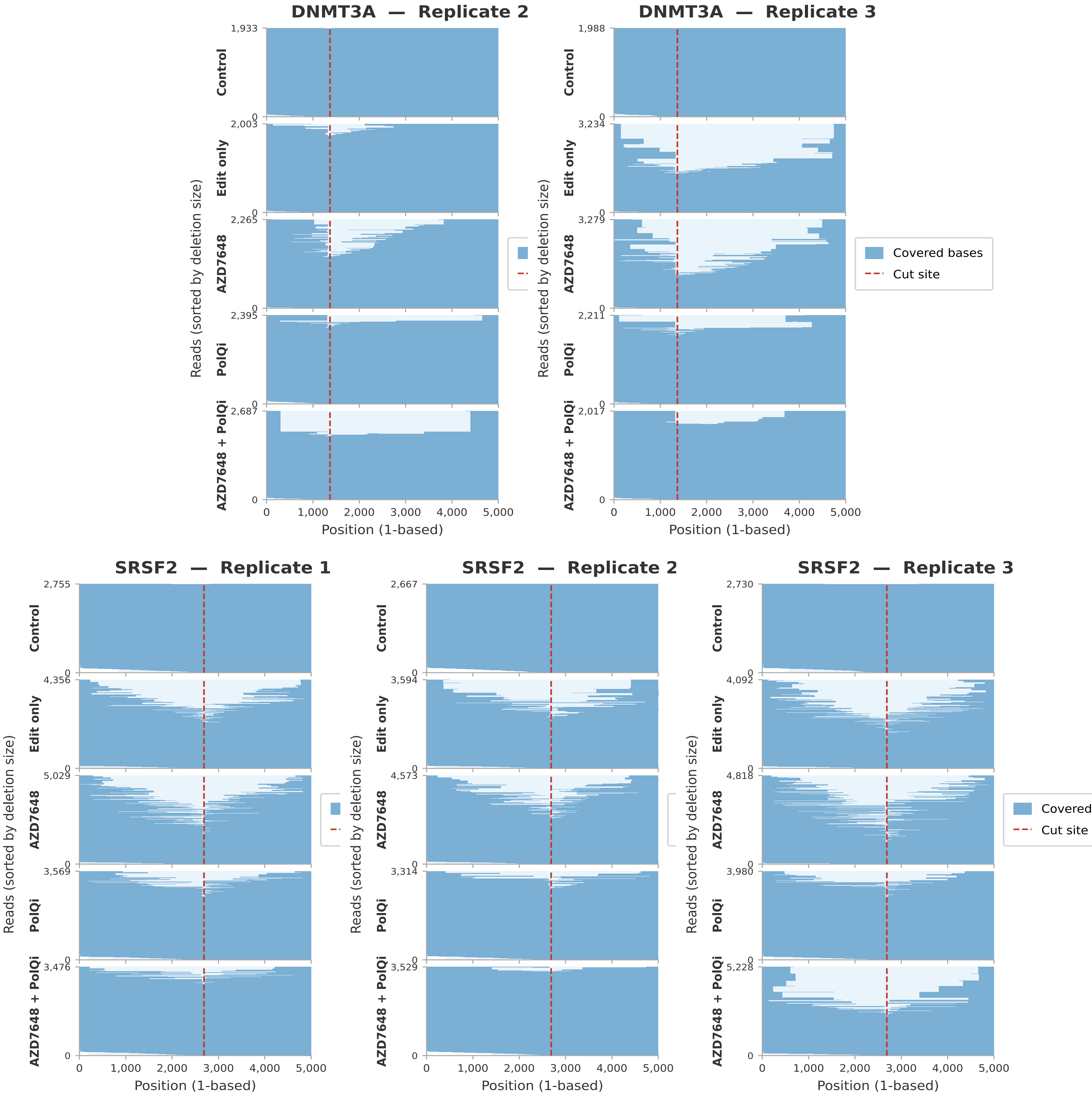


**Supplementary Figure 2. Per read coverage panels for each replicate of batch 1 for large deletion assessment.** Representative deletion size distributions, sorted by large deletion size, from Nanopore long-read sequencing at the DNMT3A loci for replicate 2 and 3 and at the SRSF2 for replicate 1, 2 and 3 detected across samples. Only reads anchored on both sides of the cut site (bilateral anchoring) are included in the analysis. Anchored reads are shown in blue, cas9 target site as a red dashed line, number of reads per condition is indicated on the top left Y axis.

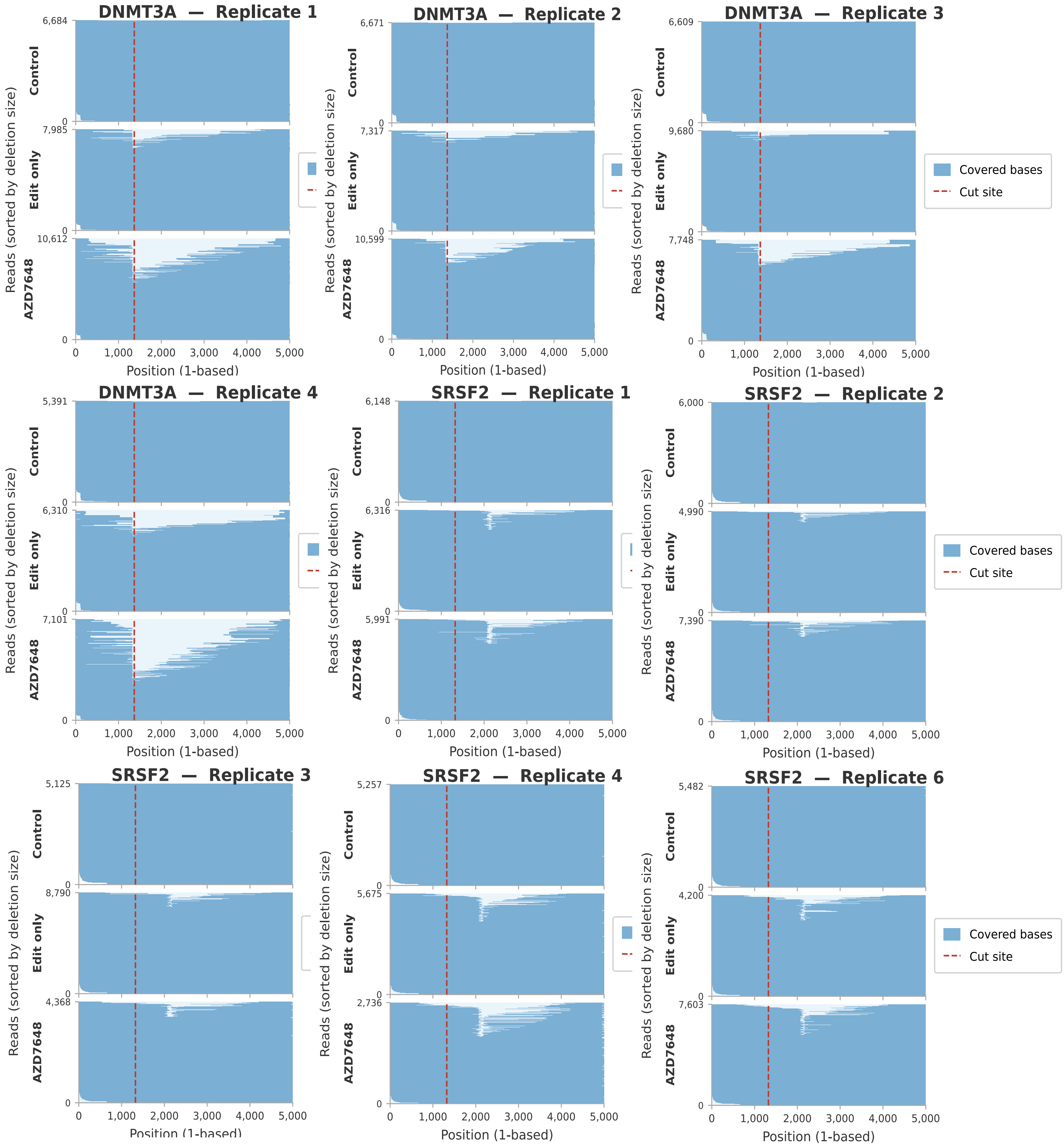

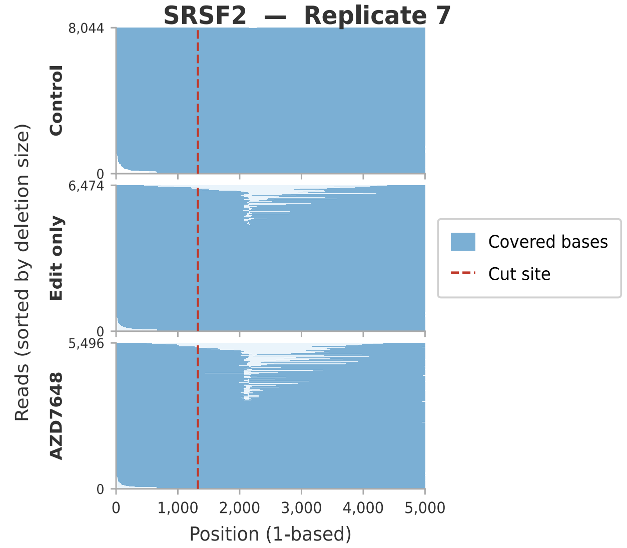


**Supplementary Figure 3. Per read coverage panels for each replicate of batch 2 for large deletion assessment.** Representative deletion size distributions, sorted by large deletion size, from Nanopore long-read sequencing at the DNMT3A loci for replicate 1, 2, 3 and 4 (Replicates 5, 6 and 7 have been discarded because of having less than a 1000 reads sequenced for the Control condition) and at the SRSF2 for replicate 1, 2, 3, 4, 6 and 7 (Replicate 5 has been discarded because of having less than a 1000 reads sequenced for the Control condition) detected across samples. Only reads anchored on both sides of the cut site (bilateral anchoring) are included in the analysis. Anchored reads are shown in blue, cas9 target site as a red dashed line, number of reads per condition is indicated on the top left Y axis.

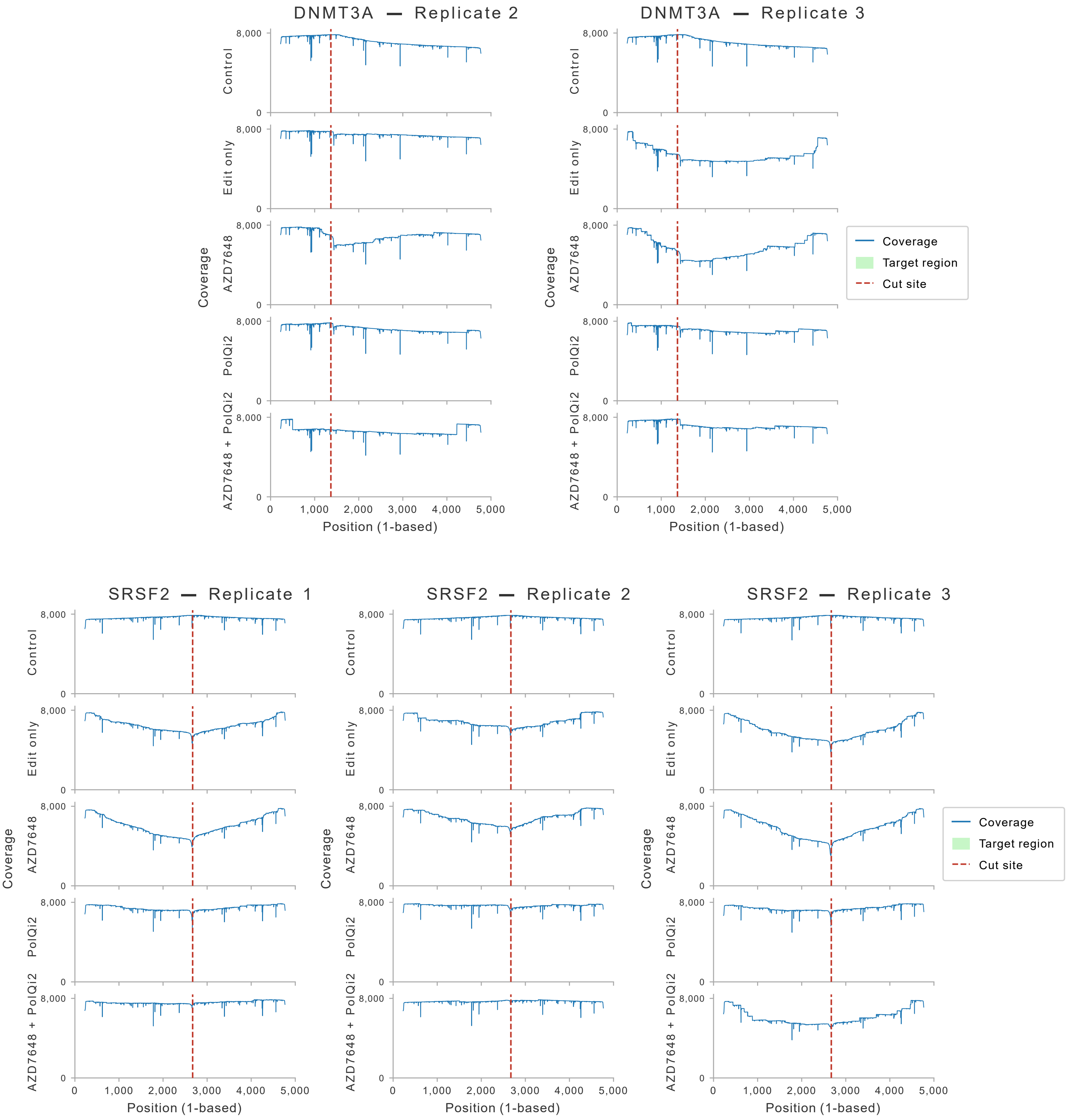


**Supplementary Figure 4. Coverage profile for all replicates from batch 1.** Coverage profile across the PCR amplicon at a representative locus, illustrating the drop in read coverage toward the 3' end characteristic of truncated Nanopore reads for both locus SRSF2 and DMT3A. These truncated reads, which cannot be assigned a true deletion size, are removed prior to large deletion quantification by the bilateral anchoring filter, ensuring that coverage drop-off artifacts do not inflate large deletion estimates.

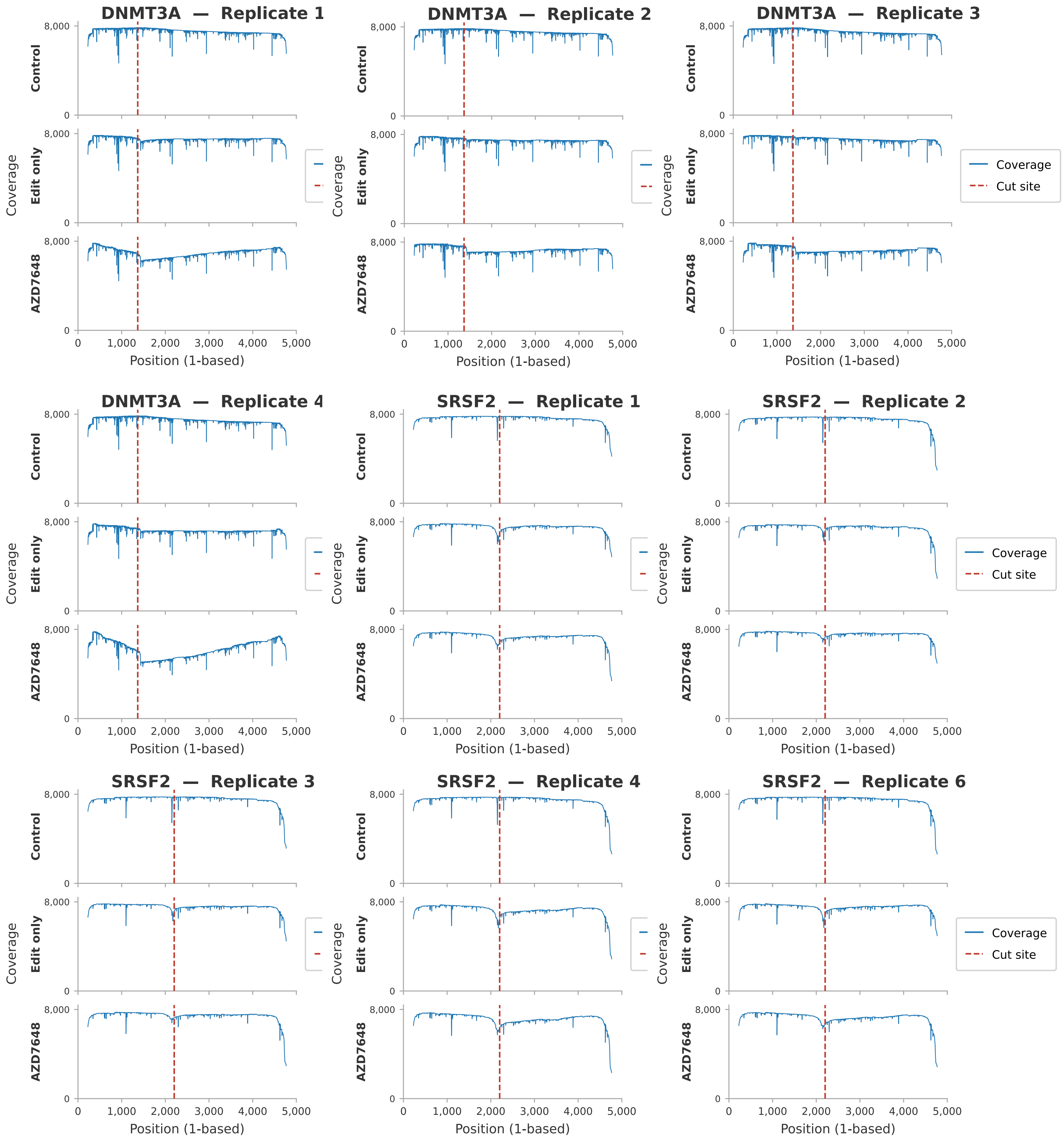


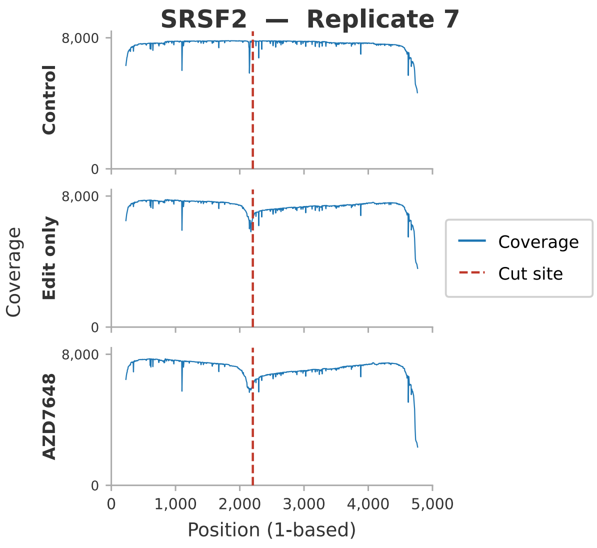


**Supplementary Figure 5. Coverage profile for all replicates from batch 2.** Coverage profile across the PCR amplicon at a representative locus, illustrating the drop in read coverage toward the 3' end characteristic of truncated Nanopore reads for both locus SRSF2 and DMT3A. These truncated reads, which cannot be assigned a true deletion size, are removed prior to large deletion quantification by the bilateral anchoring filter, ensuring that coverage artifacts do not inflate large deletion estimates.

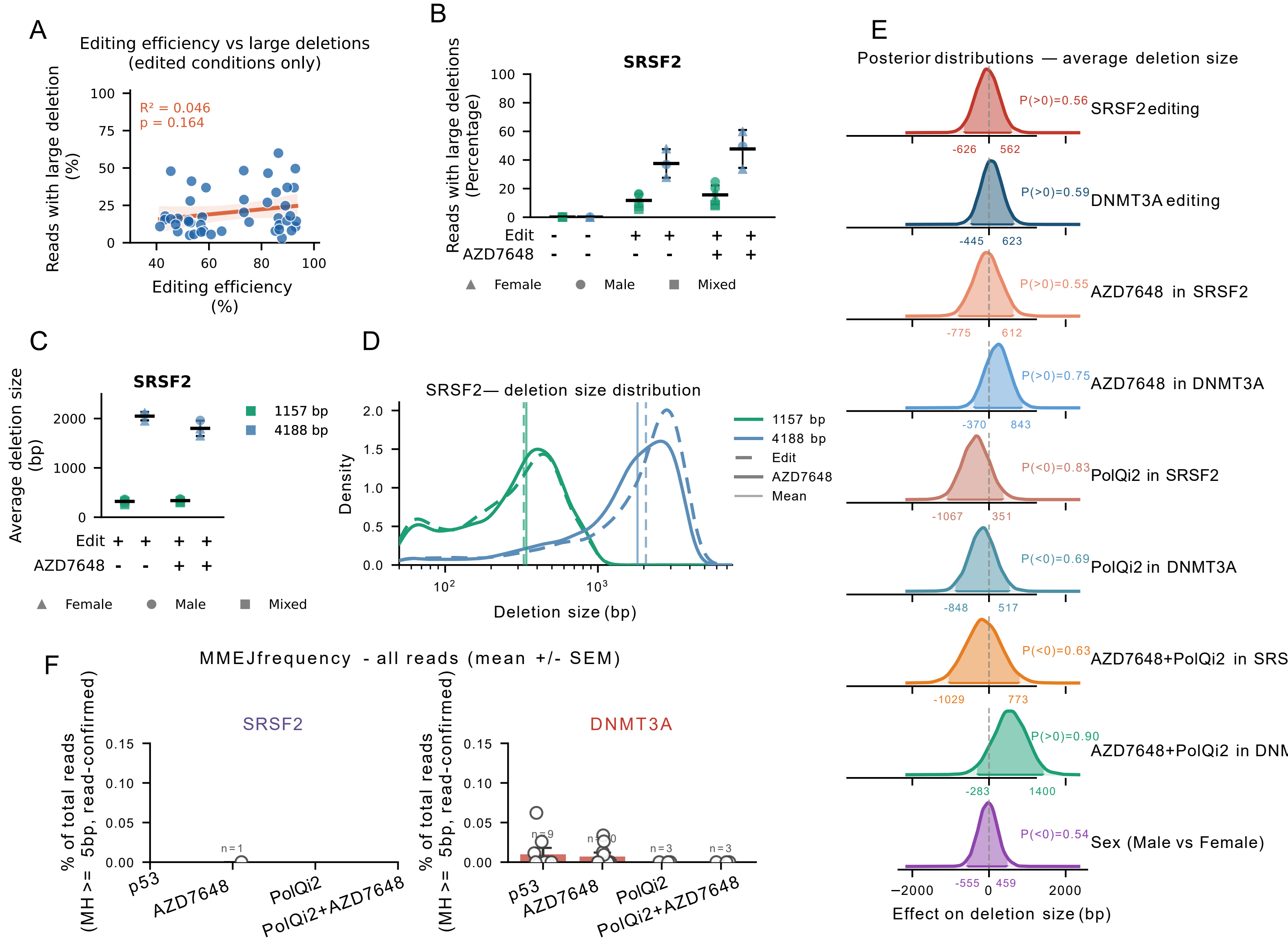


**Supplementary Figure 6. Large deletion characteristics at CRISPR-edited loci: amplicon size effects, repair pathway contributions, and editing efficiency independence. (A)** Correlation between Nanopore-derived editing efficiency and the percentage of reads carrying large deletions across edited conditions (Ctrl excluded). Each data point represents one replicate × condition × locus measurement. The solid line shows the OLS regression fit; the shaded band represents the 95% confidence interval. The absence of correlation ($R^2 = 0.021$, $p = 0.353$) indicates that large deletion frequency is not a byproduct of editing rate but is determined by the specific repair pathway context imposed by each experimental condition. **(B)** Comparison of raw large-deletion frequencies detected with two SRSF2 PCR amplicons of different sizes (1157 bp and 4188 bp) across Ctrl, no molecule, and AZD conditions. Within each condition, the two amplicon sizes are plotted side by side. Color encodes amplicon size (teal = 1157 bp; blue = 4188 bp); marker shape encodes donor sex (triangle = female, circle = male, square = mixed). The 4188 bp amplicon consistently yields higher detected large deletion frequencies, reflecting the greater sequence space available for deletion detection in longer amplicons. The two amplicons should not be compared quantitatively across batches. **(C-D)** Average size (bp) of detected large deletions for the two SRSF2 amplicons (1157 bp and 4188 bp) under no molecule and AZD conditions. The longer amplicon captures larger average deletion sizes, consistent with the interpretation that the 1157 bp amplicon truncates the detectable deletion size range and systematically underestimates both the frequency and magnitude of large deletion events. These data support the use of amplicons of at least ~4 kb for accurate large deletion quantification at CRISPR-edited loci. **(E)** Posterior probability density distributions for mixed-effect parameters from the Bayesian mixed-effects Gaussian linear model of average large deletion size (bp). The model formula was identical to (Fig1C): *avg_del_size ~ (SRSF2 + DNMT3A) × AZD7648 × PolQi2 + Sex + (1 | Replicate)*, with an identity link function and a Gaussian likelihood. Fixed and random effects specifications and reference levels were identical to the frequency model. Weakly informative priors were specified on the base pair scale: normal(0, 500) for fixed effects, Student-t(3, 1500, 500) for the intercept (centred near the observed mean of 1,482 bp), half-Student-t(3, 0, 500) for the residual standard deviation, and half-normal(0, 300) for random effect standard deviations. Parameters shown are composite contrasts identical in structure to (Fig. 1C), displayed directly in base pairs (no transformation); a dashed vertical reference line is shown at 0 bp (null effect). Positive values indicate larger deletions; negative values indicate smaller deletions. Shaded regions and HDI annotations as in (Fig. 1C). No fixed-effect parameter reached a posterior probability > 0.95, indicating that experimental conditions modulate the frequency rather than the characteristic size of large deletions. **(F)** Frequency of CRISPR-induced deletion events bearing read-confirmed microhomology ≥ 5 bp at the DNMT3A R882 and SRSF2 P95 cut site junctions, expressed as percentage of total reads. Analysis was performed on all Nanopore reads across two independent experiments (no molecule and AZD7648: n = 9 replicates; PolQi2 and PolQi2+AZD7648: n = 3 replicates). Each dot represents one biological replicate; marker shape indicates donor sex (triangle = female, circle = male, square = mixed); horizontal bar = mean; error bars = ±1 SD.

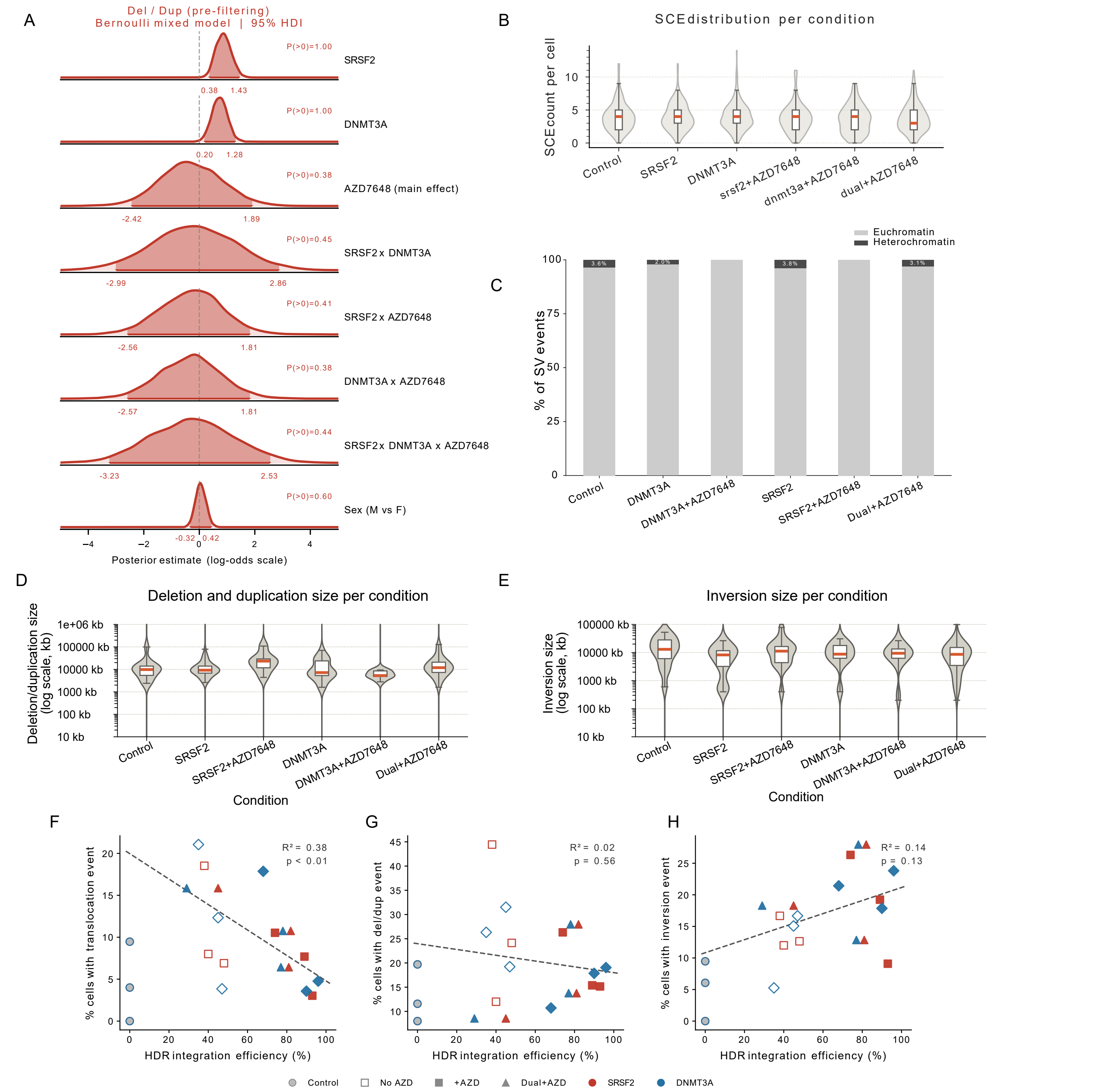


**Supplementary Figure 7. AZD7648 does not substantially alter genome-wide structural variant burden, sister chromatid exchange frequency, or SV size distributions in CRISPR-edited HSPCs. (A)** Bayesian mixed-effects analysis of del/dup cell-positive rates under the initial 5 Mb size threshold (pre-filtering sensitivity analysis). Posterior density plots show the effect of each experimental contrast on the log-odds of a cell carrying at least one deletion or duplication event of ≥5 Mb, prior to application of the arm-weakness filter and chromosome-arm size criterion. Layout and encoding are identical to panel B. This analysis confirms that the elevated del/dup burden in SRSF2-edited (P(FC>1) = 1.000) and DNMT3A-edited cells (P(FC>1) = 0.997) is robust to the choice of size threshold, while the modest AZD7648 reduction trend observed at this threshold (P(FC<1) = 0.616) becomes less apparent after applying more stringent filters, consistent with the 5 Mb threshold retaining a higher proportion of small recurrent technical events. **(B)** Per-cell sister chromatid exchange (SCE) counts across all conditions. Each point represents one single cell; the distribution of SCE counts per condition is shown across all 961 cells from three biological replicates. SCE events were extracted from the MosaiCatcher stringent quality filters (minimum allele frequency = 5%, minimum inter-SCE distance = 500 kb). Conditions are shown on the x-axis (Ctrl, SRSF2, SRSF2 + AZD, DNMT3A, DNMT3A + AZD, Dual + AZD). **(C)** Chromatin region distribution of SV events. Stacked bar plot showing the proportion of validated SV events (del/dup and inversions combined) occurring in euchromatin or heterochromatin regions, per condition. Colours: euchromatin = light grey, heterochromatin = dark grey. **(D)** Deletion and duplication size distribution per condition after filtering steps. Violin plots show the full distribution; overlaid boxplots show median (red line) and interquartile range. Y-axis in log scale kilobases (kb). No significant differences in event size were detected between conditions (Mann-Whitney U test). **(E)** Inversion distribution per condition after filtering steps. Violin plots show the full distribution; overlaid boxplots show median (red line) and interquartile range. Y-axis in log scale kilobases (kb). No significant differences in event size were detected between conditions (Mann-Whitney U test). **(F)** Correlation between HDR integration efficiency (%) and the percentage of cells carrying at least one validated reciprocal translocation event across all edited conditions and biological replicates. Each data point represents one condition × replicate × locus combination; colour indicates the edited locus (red = SRSF2, blue = DNMT3A); filled markers indicate conditions with AZD7648, open markers indicate conditions without AZD7648; triangles indicate Dual+AZD conditions; circles indicate Control. The dashed line shows the ordinary least-squares regression fit across all non-control data points. $R^2$ and p-value are indicated. **(G)** Correlation between HDR integration efficiency (%) and the percentage of cells carrying at least one validated deletion or duplication event (stringent filter: ≥10% chromosome arm length with arm-weakness filter applied). Visual encoding as in (F). **(H)** Correlation between HDR integration efficiency (%) and the percentage of cells carrying at least one validated inversion event (arm-weakness filter with shared-control blacklist at reciprocal overlap ≥0.6). Visual encoding as in **(F)**.

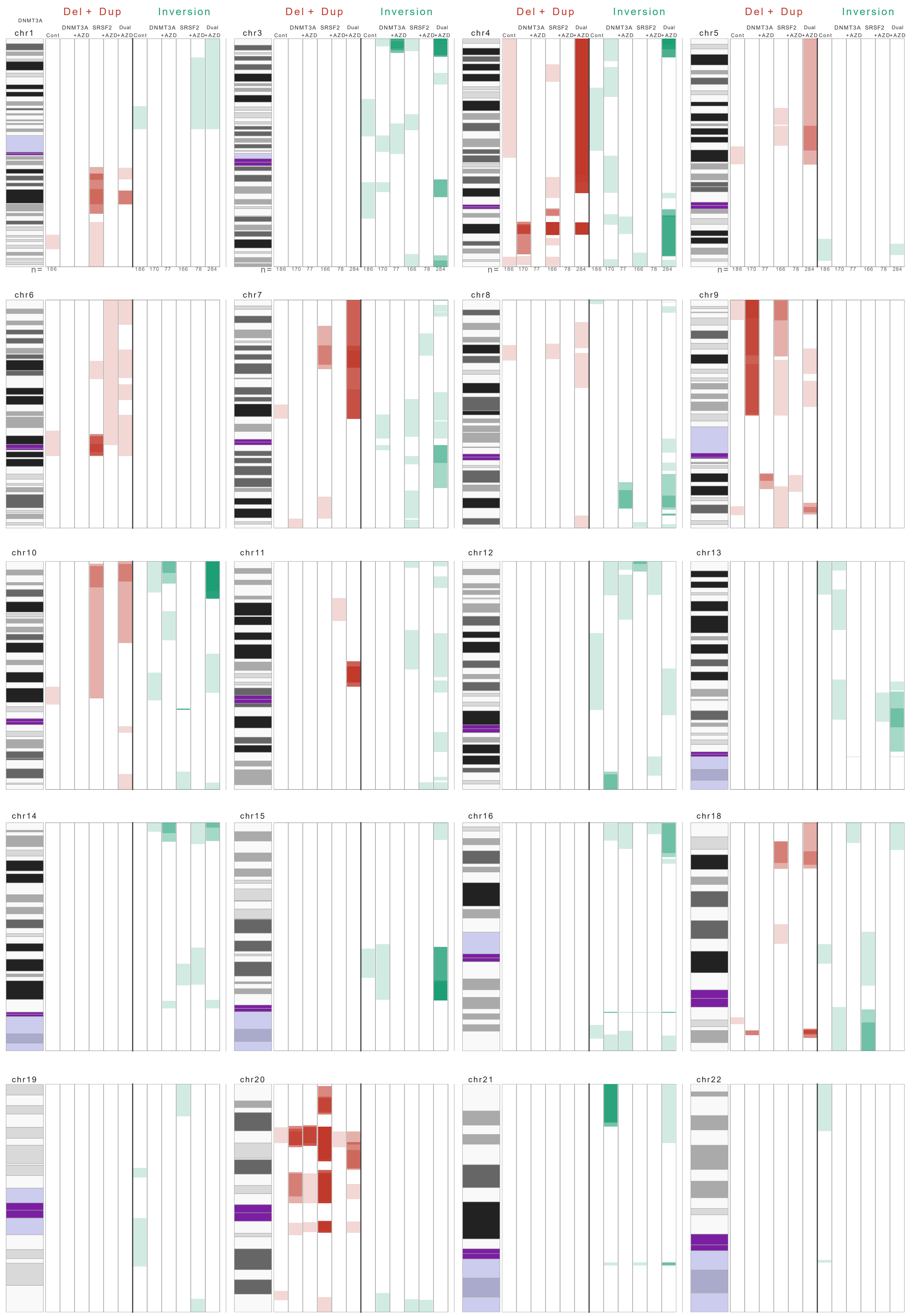


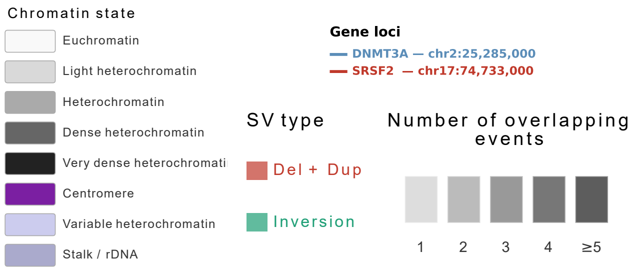


**Supplementary Figure 8. Genome-wide somatic structural variant hotspot map.** Same representation as Figure 5A for all autosomes except chr2 and chr17 (chr1, chr3–chr16, chr18–chr22). Chromosomes are arranged in groups of four per row across five rows, with all six experimental conditions displayed side by side within each chromosome group to facilitate direct comparison across conditions. For each condition, deletions and duplications combined (red) and inversions (teal) are shown as individual full-span bars to the right of the shared ideogram, with colour opacity reflecting the number of locally overlapping events. Only large structural variants representing at least 10% of the corresponding chromosome arm length were retained, and events overlapping centromeric and pericentromeric regions were excluded prior to visualization. Colour opacity reflects the number of locally overlapping events, with each additional overlapping event contributing 0.20 opacity (1 event = lightest, ≥5 events = near opaque).
