## Supplementary material for "Genome-wide structural variant landscape following HDR-enhanced CRISPR editing in human hematopoietic stem and progenitor cells": Key resources table

| REAGENT or RESOURCE | SOURCE | IDENTIFIER |
| --- | --- | --- |
| Antibodies | | |
| AF647 Mouse Anti-Human CD34 (clone 581) | Cedarlane | 343508 |
| V450 Mouse Anti-Human CD45RA (clone HI100) | BD Biosciences | 560362 |
| PE-CF594 Mouse Anti-Human CD90 (clone 5E10) | BD Biosciences | 562385 |
| CD49c Antibody, anti-human, REAfinity FITC) | Miltenyi | 130-105-364 |
| Bacterial and virus strains | | |
| Biological samples |  |  |
| Human Cord Blood for CD34+ cells harvest | Héma Québec via St Justine hospital |  |
| Chemicals, peptides, and recombinant proteins | | |
| Alt-R® S.p. Cas9 Nuclease V3, 500 µg | IDT DNA | 1081058 |
| Proteinase K, Molecular Biology Grade | NEB | P8107S |
| Alt-R® Cas9 Electroporation Enhancer, 2 nmol | IDT DNA | 1075915 |
| PLATINUM SUPERFI II MASTER MIX | Life Technologies | 12368050 |
| UM 171 | ExcellThera |  |
| AZD 7648 | Cedarlane (Cayman) | 28598-1 |
| p53 siRNA id s605 | Thermo | 4390824 |
| DIMETHYL SULFOXIDE (DMSO), Sterile | BioShop | DMS666.100 |
| P3 Primary Cell 4D-Nucleofector® X Kit L | Lonza | V4XP-3024 |
| SCF, Human (P. pastoris-expressed) | Cedarlane (GeneScript) | Z02692-10 |
| IL-3, Human(CHO-expressed) | Cedarlane (GeneScript) | Z02991-10 |
| Flt-3L, Human | Cedarlane (GeneScript) | Z02926-10 |
| IL-6, Human(CHO-expressed) | Cedarlane (GeneScript) | Z03134-50 |
| GM-CSF, Human(CHO-expressed) | Cedarlane (GeneScript) | Z02983-10 |
| Critical commercial assays | | |
| EasySep™ Human CD34 Positive Selection Kit II | STEMCELL Technologies | 17896 |
| Deposited data | | |
| Experimental models: Cell lines | | |
| Experimental models: Organisms/strains | | |
| Oligonucleotides | | |
| Alt-R® CRISPR-Cas9 tracrRNA | IDT DNA | 1072533 |
| Alt-R® CRISPR-Cas9 crRNA | IDT DNA | Custom |
| Primer 0077 R | IDT DNA | Custom |
| Primer 0077 F | IDT DNA | Custom |
| Primer 0003-A1 | IDT DNA | Custom |
| Primer 0503 F | IDT DNA | Custom |
| Primer 0503 R | IDT DNA | Custom |
| Primer 0245 F | IDT DNA | Custom |
| Primer 0245 R | IDT DNA | Custom |
| Primer 0374 F | IDT DNA | Custom |
| Primer 0375 R | IDT DNA | Custom |
| Recombinant DNA | | |
| Software and algorithms | | |
| Synthego Performance Analysis V3 | ICE Analysis | https://www.synthego.com |
| R (version 4.4.3) | R Core Team | https://www.r-project.org/ |
| Other | | |
| FBS Canadien | Thermo | 12483020 |
| RPMI1640 | LifeTech | 11875119 |
| StemSpan™ SFEM II | STEMCELL Technologies | 9655 |

| Name | Sequence | Length | Company |
| --- | --- | --- | --- |
| SRSF2 : Alt-R® CRISPR-Cas9 crRNA | CGGCUGUGGUGUGAGUCCGG | 20bp | IDT DNA |
| DNMT3A : Alt-R® CRISPR-Cas9 crRNA | GCAGUCUCUGCCUCGCCAAG | 19bp | IDT DNA |
| Primer 0077-R | TCGCGACCTGGATTTGGATT | 20bp | IDT DNA |
| Primer 0077-F | AGCGATATAAACGGGCGCAG | 22bp | IDT DNA |
| Primer 0503-F | GGCACTGAGAAGAGAAAATGCC | 22bp | IDT DNA |
| Primer 0503-R | ATCTGAAGTCGTTCACCTCACT | 20bp | IDT DNA |
| Primer 0003-A1 | TCGGCGACGTGTACATCC | 18bp | IDT DNA |
| Primer 0245-F | GCAGAACTAAGCAGGCGTCA | 20bp | IDT DNA |
| Primer 0245-R | GGCAAAGCCCTCCGGTATT | 19bp | IDT DNA |
| Primer 0374-F | CCAGGTGGGGTTTTGACTGT | 20bp | IDT DNA |
| Primer 0375-R | GTGTCTCCCGACAGACCTTG | 20bp | IDT DNA |

DNA donor templates:

- P95H ssODN :

tggacggccgcgagctgcgggtgcaaatggcgcgctacggccgcc**AT**cc**A**ga**T**tcacaccacagccgccggggaccgccaccccgcaggt

- DNMT3A ssODN :

tgactggcacgctccatgaccggcccagcagtctctgcctcgcca**GC**c**T**gctcatgttggagacgtcagtatagtggactgggaaaccaa

**Legend:**

**Silent mutation**

Guide RNA complementary sequence
